# A K_Ca_ 2.2/2.3 Opener Reverses ET-1-induced NLRP3 Activation in Hypertensive Mice Corpora Cavernosa

**DOI:** 10.1101/2024.09.06.611748

**Authors:** RS Fais, SG Comerma-Steffensen, E Pinilla, VV Matchkov, RC Tostes, FS Carneiro, U Simonsen

**Affiliations:** Department of Pharmacology, Ribeirao Preto Medical School, University of Sao Paulo, Ribeirao Preto-SP, Brazil; Department of Biomedicine, Pulmonary and Cardiovascular Pharmacology, Aarhus University, Aarhus, Denmark; Division of Pulmonary, Critical Care, and Sleep Medicine, National Jewish Health, Denver-CO, USA; Department of Biomedical Sciences/Animal Physiology, Faculty of Veterinary, Central University of Venezuela, Venezuela

**Keywords:** Erectile dysfunction, NLRP3 inflammasome, K_Ca_ 2.2/2.3, arterial hypertension

## Abstract

Hypertension-induced erectile dysfunction is associated with endothelial dysfunction in the corpus cavernosum. Membrane depolarization activates the NLRP3 inflammasome, with downregulation of endothelial Ca^2+^-activated K^+^ channels type 2.3 (K_Ca_ 2.3) and upregulation of endothelin-1 (ET-1) linked to erectile dysfunction. However, underlying mechanisms remain incompletely understood. We hypothesized that activating K_Ca_ 2.2/2.3 channels reverses erectile dysfunction and ET-1-induced NLRP3 activation in hypertensive DOCA/salt mice. Hypertension was induced in mice using a DOCA/salt model, with unilaterally nephrectomized mice as controls. We measured blood pressure, intracavernous pressure (ICP), and corpus cavernosum (CC) contractility, and performed immunoblots for K_Ca_ 2.3, caspase-1, and interleukin-1β (IL-1β). DOCA/salt mice showed impaired erectile function and increased IL-1β activity and K_Ca_ 2.3 expression. Treatment with the endothelin receptor antagonist bosentan or the K_Ca_ 2.2/2.3 channel opener NS13001 reversed these dysfunctions and reduced ET-1-induced NLRP3 activation. NS13001 also restored decreased currents in endothelial cells exposed to ET-1. These findings establish that hypertension-induced erectile dysfunction involves an ET-1/membrane depolarization/NLRP3 inflammasome axis in corpus cavernosum endothelial cells, and that targeting endothelial K_Ca_2.2/2.3 channels represents a promising therapeutic strategy to counteract erectile dysfunction.

Graphic abstract **Overview of the K_Ca_2.2/2.3 regulation on the ET-1-induced NLRP3 inflammasome activation in ECs.** NLRP3 inflammasome activation in ECs depends on endothelin receptor B. On activation, NLRP3 recruits and forms a complex with ASC as well as procaspase 1. In the final step, the assembled inflammasome platform cleaves pro-caspase-1, and caspase-1 cleaves pro–IL-1β to activate IL-1β. NS13001 activates K_Ca_2.2/2.3, which inhibits ET-1-induced NLRP3 activation. Apamin inhibits K_Ca_2.2/2.3 opening. Bosentan directly inhibits ETB receptors in ECs, preventing the NLRP3 inflammasome activation.

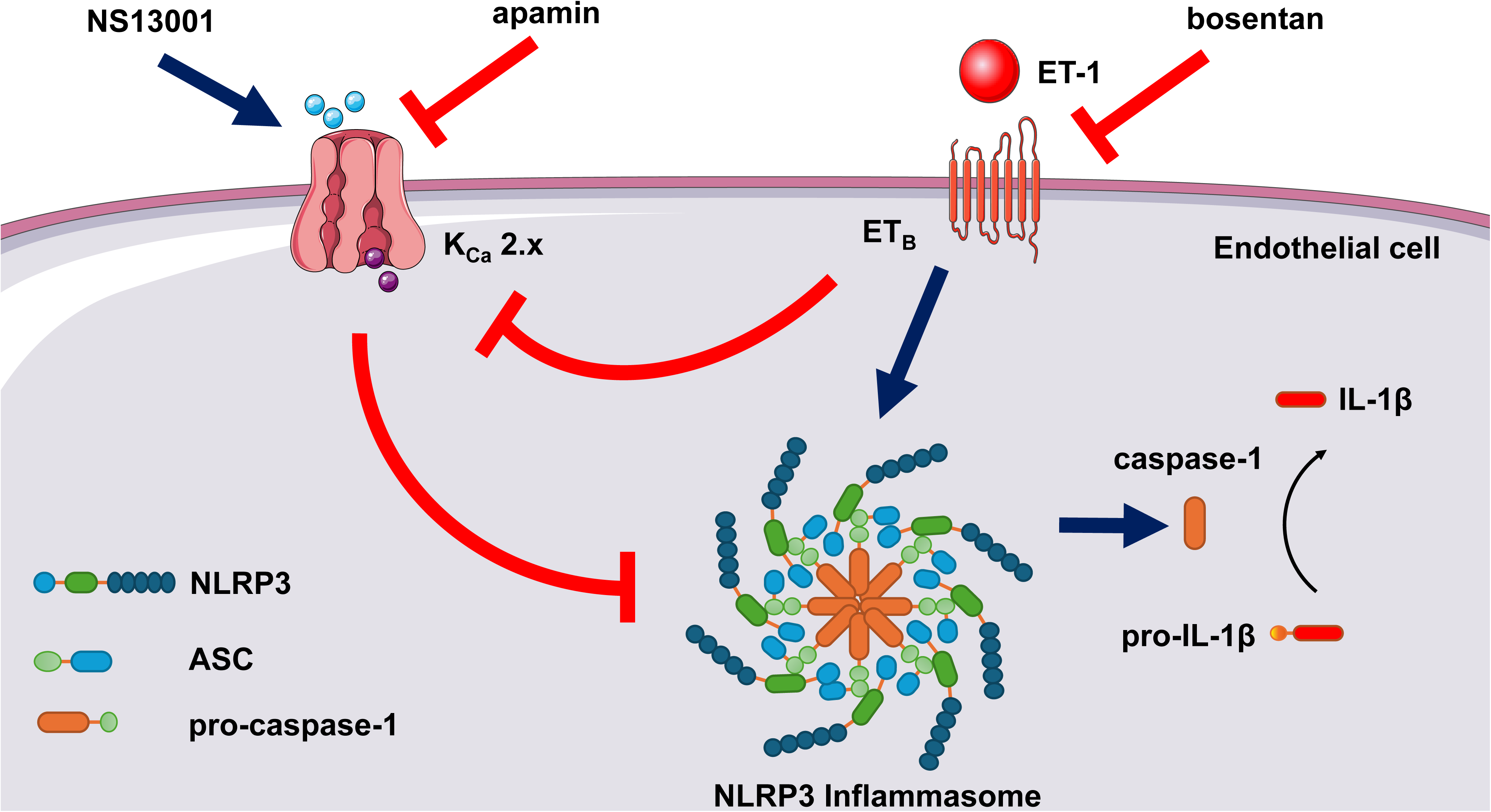

## 1. Introduction

Erectile dysfunction (ED) commonly accompanies arterial hypertension and reflects systemic endothelial dysfunction. Penile erection is mediated by endothelium-dependent vasodilation within the corpus cavernosum (CC), which enhances cavernosal blood inflow and increases sinusoidal compliance [1–5]. A key endothelial mechanism underlying this response is membrane hyperpolarization mediated by small-conductance Ca^2+^-activated K^+^ channels (K_Ca_2.x), which amplifies Ca^2+^ entry, sustains endothelial nitric oxide synthase (eNOS) activity, and promotes the release of vasodilators that relax cavernosal smooth muscle [6–8]. The K_Ca_ is a diverse group of potassium channels and are divided into small SK_Ca_ (K_Ca_2.1, K_Ca_2.2, and, K_Ca_2.3), intermediate (IK_Ca_ or (K_Ca_3.1), and large conductance Ca^2+^-activated K^+^ channels (BK_Ca_ or K_Ca_1.1) [9].

ED is a predictor of cardiovascular disease, stroke, and perhaps all-cause mortality – through an augmentation in cardiovascular mortality and independently of traditional risk factors[10,11].The K_Ca_2.3 channel is essential for nitric oxide (NO) signaling through endothelial NO synthase (eNOS) activation. NO produced by eNOS in the vascular bed not only mediates vasodilatation but also has anti-inflammatory functions [12,13]. The downregulation of K_Ca_2.3 leads to endothelial dysfunction and ED, and expression and/or function of K_Ca_2.3 channels are reduced in arterial hypertension. [6,14]

The activation of K_Ca_2.3 reduces blood pressure[3] and improves erectile function in rats. Furthermore, KCa2.3 has an anti-inflammatory effect, mainly by inhibiting the release of proinflammatory cytokines such as interleukin (IL)-6 and tumor necrosis factor-alpha (TNF-α) and by inhibiting the migration of immune cells. Thus, K_Ca_2.3 channel plays an essential role in the modulation of the immune system [12,13].

ET-1 is closely linked with endothelial dysfunction, mainly by the modulation of various populations of K^+^ channels, such as inward-rectifying (K_ir_), ATP-sensitive (K_ATP_), voltage-gated (K_v_), and K_Ca_ channels[6,15]. ET-1 has been shown to activate mononuclear phagocytes and to promote the release of TNF-α, IL-1β, and IL-6, thereby increasing the interaction between endothelial cells and monocytes. Moreover, a recent study showed that ET-1 activates NLRP3 via Ca^2+^-induced ROS generation and impairs erectile function[16–18].

Membrane depolarization is thought to be involved in the activation of NLRP3[19]. Therefore, the present study hypothesized that opening K_Ca_2.3 channels prevents the activation of NLRP3 and reverses ED. To address the hypothesis, studies were conducted on the effect of ET-1 and an opener of KCa 2.2/2.3 channels, NS13001, on the endothelial cell K^+^ currents, and the effect of NS13001 on erectile function was examined in a DOCA/salt hypertensive mouse model.

Based on these results, this study provides a novel insight into the role of K_Ca_2.3 channels in mediating the relationship between hypertension and ED, conditions that often co-exist. By revealing that ET-1-induced hypertension (DOCA/salt) impairs K_Ca_2.3 function and activates the NLRP3 inflammasome, leading to vascular and ED, it uncovers a previously underexplored critical mechanism. The therapeutic potential of the K_Ca_2.3/K_Ca_2.2 channel opener NS13001 in reversing these effects is a key finding, offering a targeted approach to treat ED and hypertension. This timely research highlights potassium channels as promising therapeutic targets for inflammatory-related vascular diseases and ED.

## 2. Materials and Methods

### 2.1. Animals

10-12 weeks-old C57BL/6 (∼25 g) male mice were housed in a room with controlled temperature (20 - 22°C) and on light/dark cycles of 12 hrs with free access to standard chow with 0.2% sodium (Altromin 1324, Brogaarden, Denmark) and filtered water. All experimental protocols adhered to the ARRIVE guidelines [20], followed Danish legislation of animal use for scientific procedures as described in the ‘Animal Testing Act’ (Consolidation Act No. 726 of 9 September 1993 as amended by Act No. 1081 of 20 December 1995) and were approved by the Danish Animal Experiments Inspectorate (permission 2019-15-0201-00009).

### 2.2. DOCA-Salt Hypertension and Arterial Blood Pressure Measurements

Deoxycorticosterone acetate (DOCA) -salt hypertension was induced as previously described [21]. Mice were anesthetized using isoflurane (5% for induction and 2% for maintenance) in 100% oxygen (2 L/min). Briefly, mice were unilaterally nephrectomized and deoxycorticosterone-acetate (DOCA; 1 g/Kg) silastic pellets were implanted subcutaneously in the scapular region. After the DOCA surgery, the mice received buprenorphine (0.05mg/kg every 8 hours for 24 hours). DOCA mice received water containing 1% NaCl and 0.2% KCl, for 3 weeks. Unilaterally nephrectomized control mice (UNI) received silastic pellets without DOCA and tap water. Animals received either the non-selective ET-1 antagonist bosentan [30 mg/day/kg (diluted in saline) of body weight, per gavage], NS13001 [1 mg/day/kg diluted in polyethylene glycol) of body weight, per gavage], apamin [1 mg/day/kg (diluted in saline) of body weight, subcutaneous], or its respective vehicle for 11 days starting 10 days after the surgery. Systolic blood pressure (SBP) was measured in mice with a tail-cuff method using a RTBP1001 blood pressure system (Kent Scientific Corporation, CT).

### 2.3. Isolation of corpus cavernosum (CC) strips and CC primary endothelial cells

For isolation of primary endothelial cells and corpus cavernosum (CC) strips, the mouse was gently placed in a transparent chamber (19 cm wide x 29 cm deep x 13 cm high) and exposed to isoflurane 3% in 100% oxygen (2L/min). Once the pain reflexes disappeared, the animals were decapitated. After this procedure, the CC was isolated and split into 2 strips, one was used for CC contractility measurements, and the other for immunoblotting.

### 2.4. Drugs and solutions

Physiological Krebs-Henseleit buffer of the following composition was used: NaCl 130 mM, KCl 4.7 mM, KH_2_PO_4_ 1.18 mM, MgSO_4_•7H_2_O 1.17 mM, NaHCO_3_ 14.9 mM, EDTA 0.026 mM, CaCl_2_•2H_2_O 1.6 mM and D-glucose 5.55 mM. A high concentration of KCl buffer of the following composition was used: NaCl 14 mM, KCl 120 mM, KH_2_PO_4_ 1.18 mM, MgSO_4_•7H_2_O 1.17 mM, NaHCO_3_ 14.9 mM, EDTA 0.026 mM, CaCl_2•_2H_2_O 1.6 mM and D-glucose 5.55 mM. pH in all buffers was adjusted to 7.35 before its use. The incubations were performed with ET-1 (100 nM, Tocris 1160; diluted in 5% BSA + 95% deionized water), MCC950 (1 µM, Cayman Chemical 17510; diluted in 5% DMSO and 95% deionized water), NS13001 (100 nM, ChemShuttle, CA, USA; diluted in polyethylene glycol), apamin (100 nM, Latoxan L8407; diluted in deionized water). The in vivo experiments were performed with bosentan (30 mg/day/kg, Sigma Aldrich 157212-55-0; diluted in saline), NS13001 (1 mg/day/kg, ChemShuttle, CA, USA; diluted in polyethylene glycol), apamin (1 mg/day/kg; Latoxan L8407; diluted in saline).

### 2.5. Culture of primary endothelial cells of Corpus cavernosum

Endothelial cells from CC of intact C57BL/6 mice were isolated as previously described[3]. Briefly, a freshly isolated CC strip was placed in sterile PBS and cut into small pieces together with collagenase II (400 mg ml−1, Worthington Biochemical Corporation, Lakewood, NJ, USA) for 60 min at 37°C. The sampled tissue was centrifuged at 900 RPM with isolation of the pellet and transferred to a coated gelatin and collagen Petri dish at 37°C. The culture media was Dulbecco’s Modified Eagle Medium with 1% Fungizone, 1% penicillin streptomycin, newborn fetal calf serum and HEPES. When reached confluence after 1–4 days, the cells were incubated with the endothelium-specific CD102 [intracellular adhesion molecule 2 (ICAM2)] rat anti-mouse (diluted 1:500, BD Biosciences 553326. RRID: AB_394784) for 30 min followed by incubation with Dynabeads with sheep anti-rat IgG (Life Technologies, Oslo, Norway) for 30 min at 4°C. Trypsin was added to separate the cells from Petri dish and transferred to dynamag 5TM (Life Technologies, Oslo, Norway) for 2 min cleaning. The pellet was dissolved and left in a coated Petri dish for further cultivation in endothelial cell growth serum (Provitro AG, Berlin, Germany). The endothelial cells were treated with ET-1 (4 h) in the presence of NS13001 (30 min), apamin (30 min), or vehicle. After the treatment, the cells were separated for patch clamp or Western blot experiments.

### 2.6. Patch-Clamp Electrophysiology

Endothelial cells were detached from Petri dish with trypsin-EDTA, washed, and patch clamped using a Port-a-Patch automated system (Nanion Technologies, Munich Germany), as previously described [3]. Briefly, 3–5 MΩ resistance microchips were used. Currents were recorded using an extracellular solution containing (in mM): 140 NaCl, 4 KCl, 10 Glucose, 10 HEPES, 2 CaCl_2_ and 1 MgCl_2_, pH 7.3–7.4. The intracellular solution was prepared as described previously (Stankevicius et al., 2011) and contained (in mM): 30 KCl, 100 K-Aspartate, 1 mM MgCl_2_, 10 mM EGTA, 3 Na_2_ATP, 5 mM HEPES, and 8,5 CaCl_2_, pH 7.2. The osmolarity of all solutions was 290–320 mOsm. All voltages were corrected for calculated junction potentials present between the internal and external solutions before seal formation. The K^+^ currents were activated by voltage steps from −140 to 140 mV in 40 mV increments for 250 ms and using a holding membrane potential of −80 mV. Currents measured at a step of 100 mV were used to compare the effects of the K_Ca_2.2/2.3 openers or blockers used in this study. All current amplitudes were expressed as current density (pA/pF) normalized to cell capacitance.

### 2.7. In vivo measurements of intracavernosal pressure and mean arterial pressure

Mice were anesthetized with 2% isoflurane for induction and maintained with 1.5% in 100% oxygen (2 L/min) [4]. Then, the left carotid artery and right CC of each mouse were cannulated for continuous monitoring of mean arterial pressure (MAP) and intracavernosal pressure (ICP), respectively.

In the DOCA and UNI mice treated with either vehicle, bosentan, NS13001, or apamin, the cavernosal nerve was electrically stimulated with silver electrodes at different frequencies (5 V, 1 ms pulses, and frequencies between 0.5 and 20 Hz for 60 sec stimulations) to induce changes in ICP. Each frequency was followed by a 2-minute resting period. Cavernosal nerve mediates the relaxation and contraction of cavernous smooth muscle, respectively, together regulating the erectile function of penile tissue(Dean and Lue, 2005).

In intact C57BL/6 mice, the cavernosal nerve was stimulated electrically with silver electrodes at 6 V, 1 ms pulse width, for 60 sec, and the frequency at 16 Hz to induce changes in ICP and obtain the maximum (max) ICP/MAP. Subsequently, the cavernosal nerve was stimulated at the same parameters and varying the frequency from 2 to 8 Hz to determine the 50% (submaximal) of ICP/MAP for each mouse(Sobrano Fais et al., 2023). Then, we administered intracavernosal NS13001 (0.1 µg/Kg), apamin (20.27 µg/Kg), or vehicle. A new submaximal stimulation was performed at 3, 10, 20, and 30 minutes, and the maximal stimulation at 25 minutes. A second intracavernosal administration with ET-1 (0.2492 µg/Kg) or vehicle was performed, and the submaximal stimulation was performed at 3, 10, 20, and 30 minutes, and the maximal stimulation at 25 minutes.

### 2.8. Cavernosal tissue contractility

Cavernosal strips from intact C57BL/6 mice were isolated and mounted in 10 mL-myograph chambers (Danish Myo Technology, Aarhus, Denmark) containing buffer continuously bubbled with a mixture of BioAir (5% CO_2_ and 19.82% O_2_ in nitrogen) and maintained at 37°C as previously described [2]. The tissues were stretched to a resting force of 2.5 mN and allowed to equilibrate for 60 min. Changes in isometric force were recorded using the PowerLab/8SP data acquisition system (Chart software, version 7; ADInstruments, Colorado Springs, CO). At the end of the equilibration period, phenylephrine (1 μM) was added to the organ bath to verify the preparations’ contractile ability.

The contraction was evaluated by cumulative concentration-response curves to phenylephrine (1 nM to 100 µM) in CC strips. Relaxation was evaluated by constructing cumulative concentration-response curves for acetylcholine (ACh) (100 pM – 3 µM; endothelium-dependent relaxation) and sodium nitroprusside (SNP) (10 pM – 100 µM; NO donor) in CC strips pre-contracted with phenylephrine (10 µM). To investigate neurogenic relaxations in CC strips, the preparations were treated with guanethidine (30 µM) and atropine (1 µM) as previously described(Simonsen et al., 2001). Then, the preparations were contracted with phenylephrine, and electrical field stimulation (0.5 to 32.0 Hz) was performed.

### 2.9. Immunoblotting

DOCA or UNI animals received either the non-selective ET-1 antagonist bosentan [30 mg/day/Kg (diluted in saline) of body weight, per gavage], NLRP3 inhibitor [10 mg/day/Kg (diluted in saline) of body weight, subcutaneous], NS13001 [1 mg/day/Kg diluted in polyethylene glycol) of body weight, per gavage], apamin [1 mg/day/kg diluted in saline) of body weight, subcutaneous], or its respective vehicle for 11 days starting 10 days after the surgery. The samples for this immunoblotting experiments were obtained from mice exclusively used for immunoblotting, thereby avoiding interference from in vivo or ex vivo stimulation.

The CC endothelial cells were obtained from health mice and treated with ET-1 (100 nM, Tocris 1160; diluted in 5% BSA + 95% deionized water), MCC950 (1 µM, Cayman Chemical 17510; diluted in 5% DMSO and 95% deionized water), NS13001 (100 nM, ChemShuttle, CA, USA; diluted in polyethylene glycol), apamin (100 nM, Latoxan L8407; diluted in deionized water).

The isolated CC from intact C57BL/6 mice were cleaned from surrounding fat tissue and then snap-frozen in liquid nitrogen for storage at -80 °C. The samples were later homogenized in a lysis buffer [50 mM Tris/HCl, 150 mM NaCl, 1% Nonidet P40, 1 mM EDTA, 1 μg/ml leupeptin, 1 μg/ml pepstatin, 1 μg/ml aprotinin, 1 mM sodium orthovanadate, 1 mM phenylmethanesulfonyl fluoride (PMSF), and 1 mM sodium fluoride] (Fais et al., 2019). Protein concentration was determined by the Lowry assay. Spectra multicolor broad range protein ladder (10 to 260 KDa) was used as protein standard. Aliquots with 60 µg of proteins were prepared and separated by electrophoresis at 100 V for 2 hours at 4° C in 4-20% Criterion TGX Stain-Free gels (bio rad) and transferred for 1 hour to a nitrocellulose membrane at 100 V at 4° C. After transfer, total protein was visualized using stain-free technology by exposing the membranes to UV light. Nonspecific binding sites of the membrane to the primary antibodies were blocked with 0.3% iBlock solution for 1 hour at room temperature. The primary antibodies described below were incubated for 12 hours at 4° C, and the secondary antibodies were incubated for 1 hour at room temperature. Protein bands visualization was obtained by chemiluminescence after ECL reaction (Amersham ECL Prime Western Blotting Detection Reagent) and image capture was performed on ImageQuant 350 gel imager (GE Healthcare, Piscata Way, NJ, USA). The densitometric quantification was performed by ImageJ® software. Membranes were stripped with Restore 2x Western Blot Stripping Buffer (Thermo) for 30 minutes at 37° C. The densitometric quantification was normalized by the total protein in the pre-stained gel. Total protein is considered a more reliable control than housekeeping proteins [22–24].

The following antibodies were used in the study: caspase-1 (diluted 1:500, Imgenex IMG-5028. RRID: AB_316670), IL-1β [diluted 1:500, NovusBio NB600-633, RRID: AB_577977], KCa2.3 (diluted 1:500, NovusBio NBP3-46424. RRID: AB_3738376), β-actin (diluted 1:1000, Cell Signalling 5125. RRID: AB_1903890. Total protein in each lane was used as an endogenous control for normalization of all proteins. Membranes were then incubated with the following secondary antibodies: goat anti-mouse IgG H&L (diluted 1:10.000, ab6789, Abcam. RRID: AB_955439) and goat anti-rabbit IgG H&L (diluted 1:10.000, ab6721, Abcam. RRID: AB_955447).

### 2.10. Statistical analysis

The declared group size is the number of mice. Statistical analysis was undertaken only for experiments where each group size was at least n=5. The Western blot, CC reactivity, and immunohistochemistry results were analyzed by one- or two-way ANOVA and followed by Tukey post-test according to each analysis. The in vivo results were analyzed by repeated-measures two-way ANOVA and followed by Tukey post-test. The models’ assumptions were investigated by inspecting Q-Q plots and data were logarithmically transformed when necessary to generate a Gaussian-distributed data set. Relaxation responses were expressed as the percentage change from pre-contraction induced by phenylephrine. Concentration-effect curves were submitted to nonlinear regression analysis using the GraphPad Prism program (GraphPad Prism 9.0; GraphPad Software Inc., San Diego, CA, USA). Relaxant potency and maximal response were expressed as pEC50 (negative logarithm of molar concentration producing 50% of the maximal response) and Emax (maximal effect produced by the agonist), respectively. Statistical analysis of the Emax and pEC50 values was performed using nonlinear regression followed by Student’s t-test or one-way ANOVA. Values of p less than 0.05 were considered statistically significant.

## 3. Results

### Effect of endothelin-1 receptor inhibition on erectile function and NLRP3 inflammasome activity in DOCA/salt model of hypertension

In vivo cavernosal nerve stimulation in mice induced a frequency-dependent increase of ICP (figure 1b and 1c). When mice were treated with the vehicle (for bosentan), ICP/MAP was reduced in Deoxycorticosterone acetate (DOCA)/salt mice at the frequencies from 4 to 20 Hz when compared with unilateral nephrectomized (UNI) mice (Figure 1c). Additionally, DOCA/salt mice had a gradual increment of the blood pressure from day 6 to 9, but then, it was kept sustainedly increased until day 21 (Figure 1d). Bosentan, a non-selective endothelin-1 (ET-1) receptor antagonist, treatment restored the ICP/MAP (Figure 1c) and systolic blood pressure (Figure 1d) in DOCA/salt mice.

**Figure 1.**
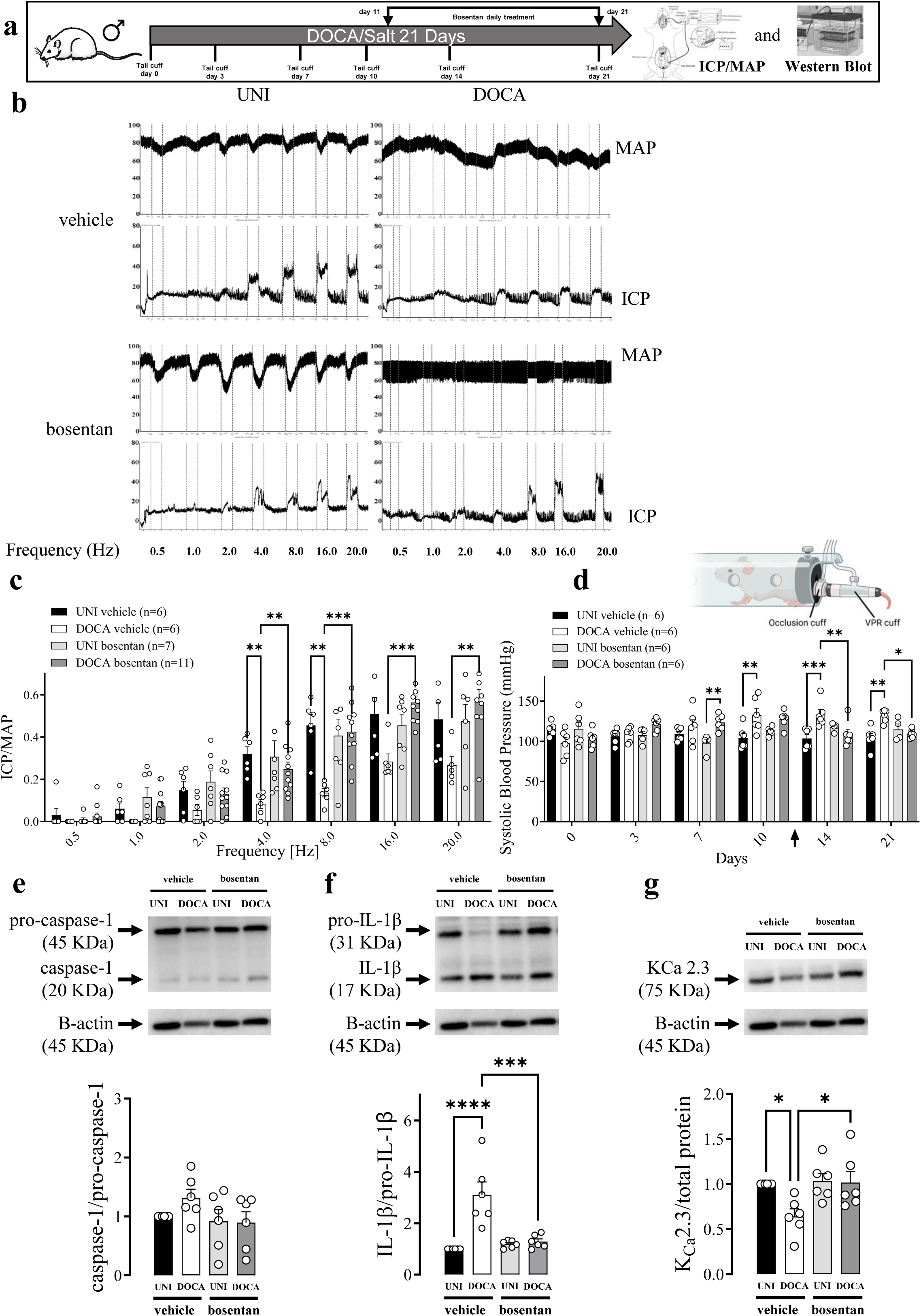
The endothelin receptor antagonist bosentan restores erectile function and inhibits inflammation in DOCA/salt mice. (**a**) Experimental design. (**b**) Original tracings showing the intracavernosal pressure (ICP, lower traces) and mean arterial pressure (MAP, upper traces) in response to cavernosal nerve stimulation assessed over a range of frequencies (0.5-20 Hz) in a unilateral nephrectomized (UNI) and a DOCA/salt (DOCA) hypertensive mice treated with vehicle or bosentan. (**c**) Average responses in ICP/MAP and (**d**) systolic blood pressure of UNI and DOCA hypertensive mice. Immunoblot analysis of (**e**) caspase-1, (**f**) IL-1β, and **(g)** K_Ca_2.3 expression in CC strips of UNI and DOCA hypertensive mice treated with bosentan or vehicle. The arrow indicates when the treatment started (c). Data represent the mean ± SEM values of the groups. * p<0.05. n=5-11. The comparison of each value was performed by two-way ANOVA followed by the Holm-Šídák post-test. ICP=intracavernosal pressure; MAP=mean arterial pressure. The entire membranes for pro-caspase-1, caspase-1, pro-IL-1β, IL-1β, and K_Ca_2.3 western blots are shown in figures S7 and S8.

In the mouse DOCA/salt model of arterial hypertension, there was a trend of increased caspase-1 activity (measured by the caspase-1/pro-caspase-1 ratio) in CC (Figure 1e). IL-1β activity (obtained by the IL-1β/pro-IL-1β) was increased in the CC from DOCA/salt mice (Figure 1f). On the other hand, K_Ca_2.3 expression was reduced in DOCA/salt mice CC. Additionally, bosentan prevented the caspase-1 activity trend, IL-1β activity increment, and K_Ca_2.3 expression reduction in CC from DOCA/salt mice (Figure 1e, 1f, and 1g). Moreover, Bosentan also prevented the reduction in the pro-IL-1β in the CC of DOCA/salt mice (Figure 1g). No significant alterations were observed in the expression of pro-caspase-1, caspase-1, and IL-1β (Figure S2a and S2b).

DOCA/salt intervention did not modify the contractility to phenylephrine or the relaxation in response to electrical field stimulation of CC (Figure 2b & 2c). Bosentan did not change phenylephrine-induced contraction or the electrical field stimulation-induced relaxation of CC (2d & 2e, Table 1). However, DOCA/salt reduced endothelium-dependent and endothelium-independent relaxations to ACh and SNP, respectively, shifting the concentration-response curves to the right (Figure 2d & 2e, Table 1). Bosentan treatment of DOCA/salt mice prevented these impairments (2d & 2e, Table 1).

**Figure 2.**
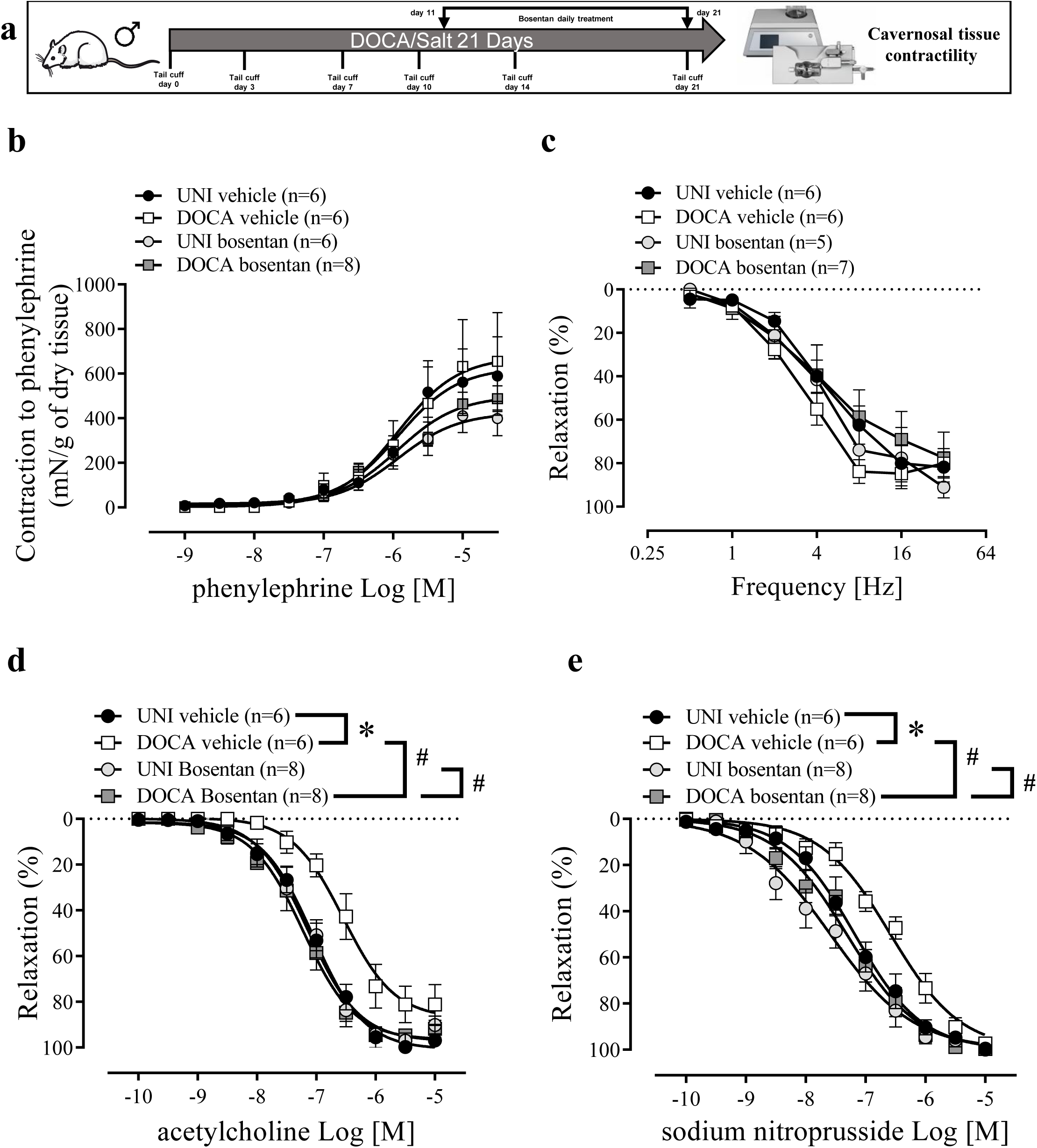
Bosentan restores the impaired endothelium-dependent and -independent relaxations in corpus cavernosum of DOCA/salt mice. (**a**) Experimental design. (**b**) Concentration-response curves for phenylephrine, (**c**) frequency-response curves for electrical-field-stimulation-induced relaxation, and concentration-response curves for (**d**) ACh and (e) SNP in CC strips of DOCA/salt (DOCA) hypertensive and unilaterally nephrectomized (UNI) mice treated with bosentan or vehicle. The values of the graphics represent mean ± SEM. * p <0.05. n = 6-8. The comparison of pEC_50_ and Emax (table 1) and each value was performed by two-way ANOVA followed by the Holm-Šídák post-test.

**Table 1.**
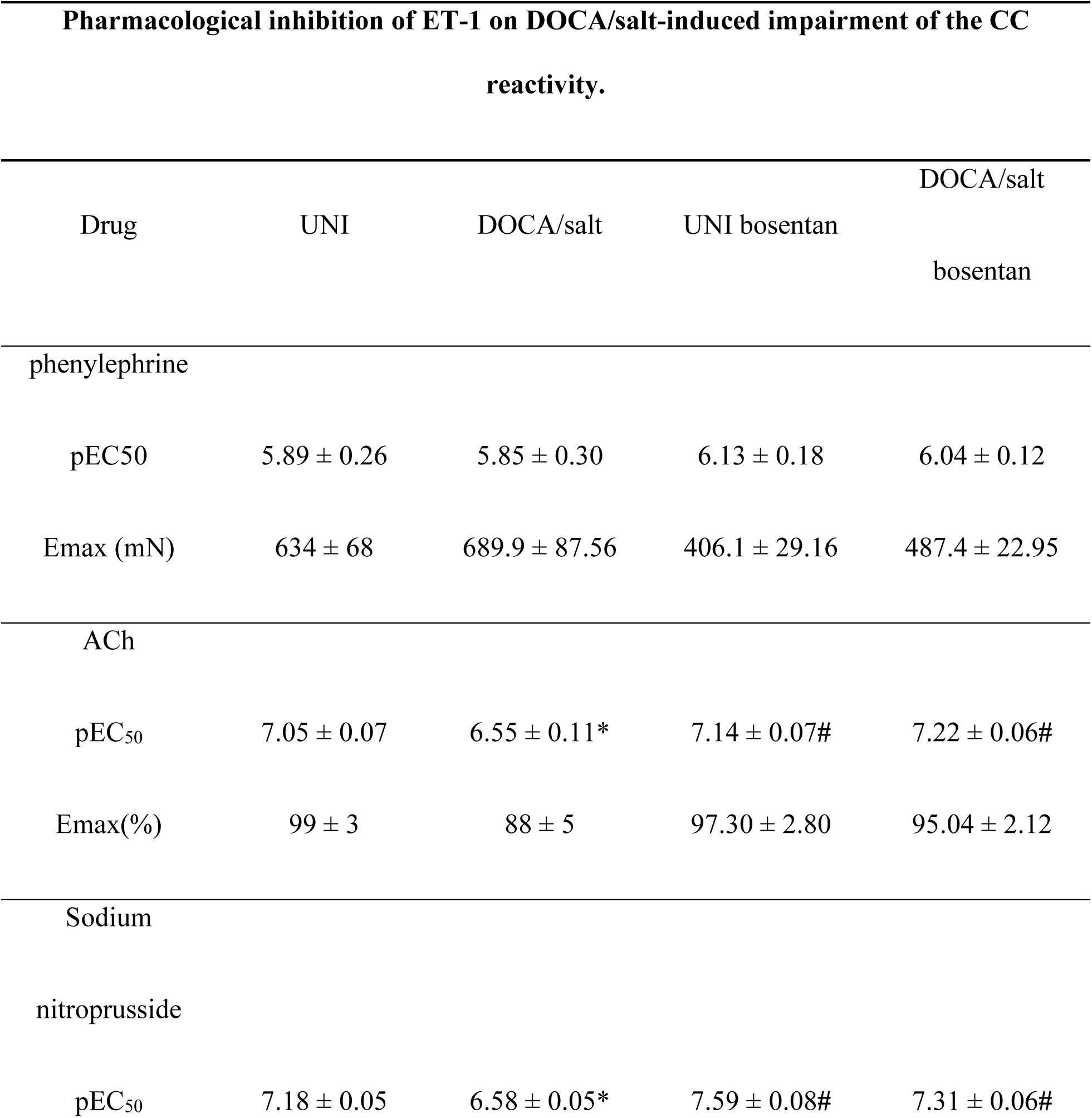
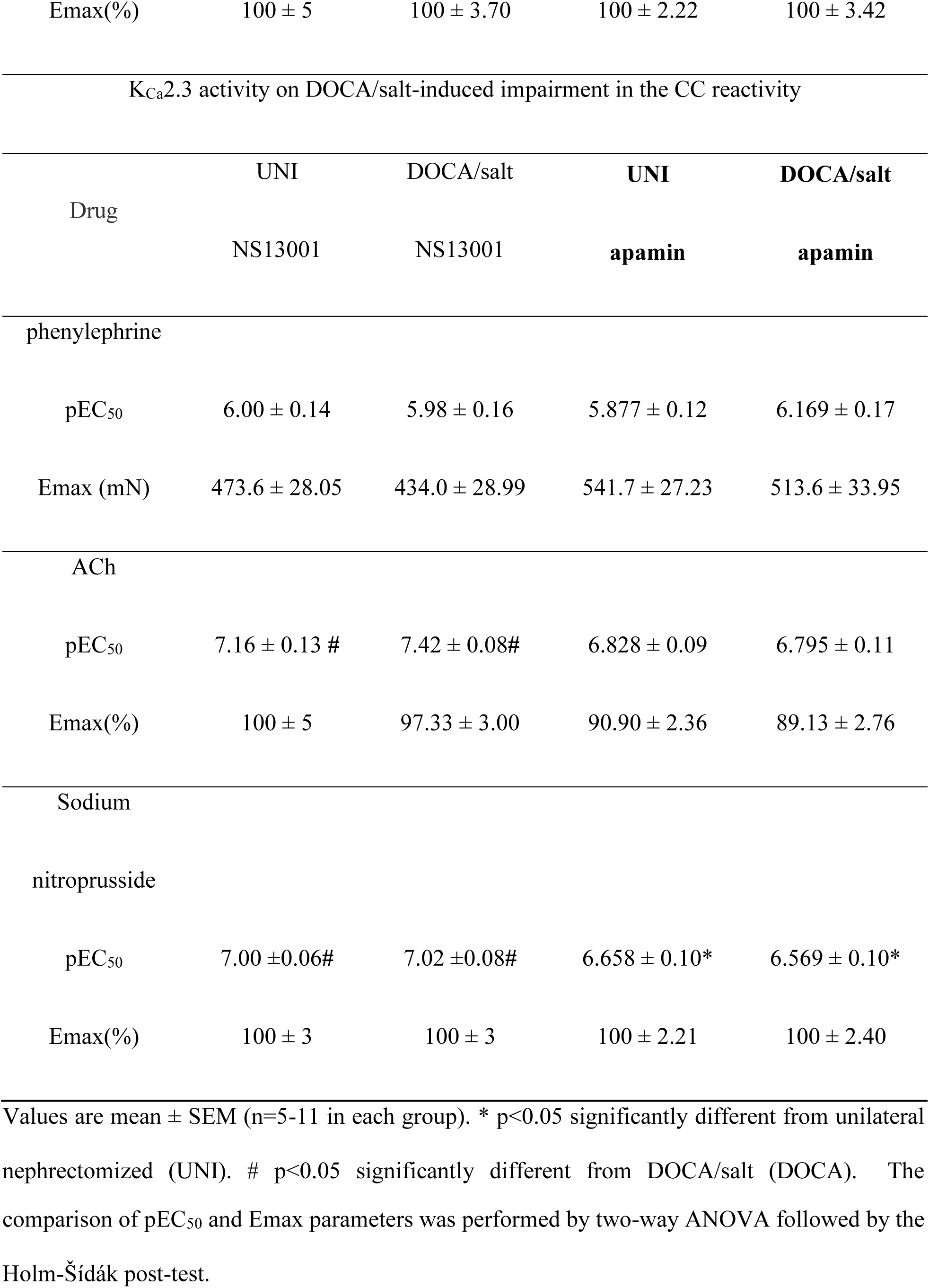
Pharmacological inhibition of ET-1 on DOCA/salt-induced impairment of the CC reactivity. Values of Emax and pEC_50_ for the phenylephrine, ACh, and SNP concentration-effect curves in CC of mice. Curves were performed in CC of unilaterally nephrectomized (UNI) or DOCA/salt-treated mice combined with the treatment with vehicle, bosentan or NS13001.

### Effect of NLRP3 inhibition on caspase-1 and IL-1β activity in DOCA/salt mice CC

In the mouse DOCA/salt model of arterial hypertension, there was an increased caspase-1 and IL-1β activity in the CC (Figure S1b). On the other hand, K_Ca_2.3 expression was reduced in DOCA/salt mice CC. Additionally, MCC950 (NLRP3 inhibitor) prevented the caspase-1 and IL-1β activity increment, and K_Ca_2.3 expression reduction in CC from DOCA/salt mice (Figure S1a, S1b and S1c). Furthermore, MCC950 also prevented the reduction in the pro-caspase-1 and pro-IL-1β in the CC of DOCA/salt mice (Figure S2c, S2d). No significant alterations were observed in the expression of caspase-1 and IL-1β (Figure S2c, S2d).

### ET-1 inhibited the KCa2.2/2.3 channels in endothelial cells, impaired the erectile function and activated the NLRP3

The activity of KCa2.2/2.3 channels using whole cell voltage clamp was measured. The K^+^ currents were measured in response to a voltage-step protocol. The total membrane conductance was significantly suppressed in the presence of ET-1 (Figure 3c). The activity of NLRP3 in endothelial cells was evaluated by Western blot for caspase-1 (Figure 3d) and IL-1β activity (Figure 3e), or K_Ca_2.3 expression. These Western blot results suggest that ET-1 administration increased the NLRP3 activity and reduced the KCa.2.3 expression.

**Figure 3.**
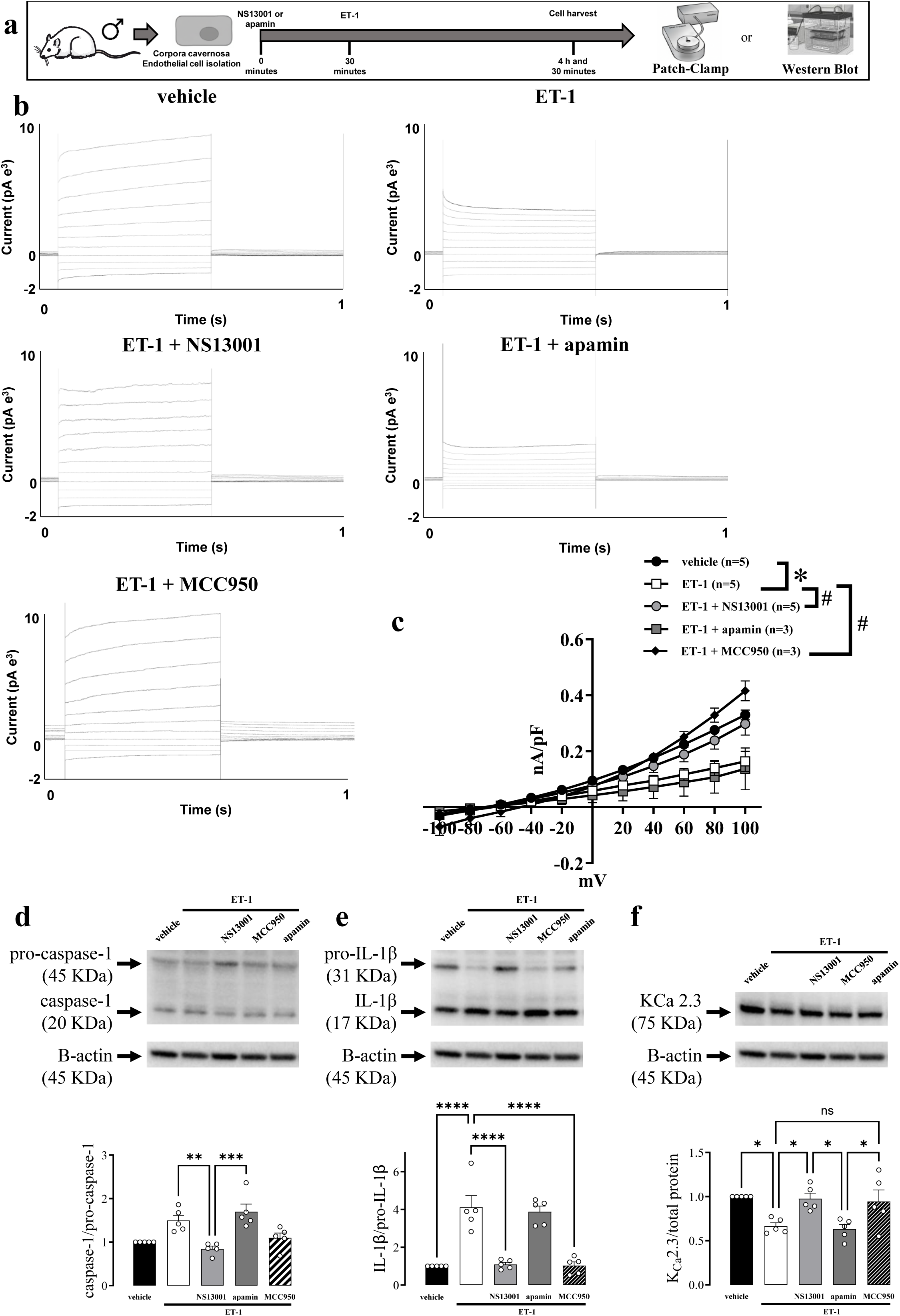
An opener of K_Ca_2.2/2.3 channels restores the background membrane conductance and inhibits inflammation in primary endothelial cells of corpus cavernosum (CC) after endothelin-1 treatment. (**a**) Experimental design. (**b**) Representative traces from the (**c**) Current evoked by the voltage-step protocol from −100 to +100 mV in endothelial cells in the presence of vehicle or ET-1 (100 nM) and the effect of NS13001 (100 nM) or apamim (100 nM) administration**. The representative traces (upper panel) and the average current-voltage relations (lower panel).** Immunoblotting (upper panel) and expression (lower panel) for (**d**) caspase-1 and (**e**) IL-1β in murine *CC* after the administration of vehicle or ET-1 alone or combined with NS13001 and apamin. The data is presented as means ± SEM. \**p* ≤ 0.05. n = 3-6. Each value was compared by two-way ANOVA followed by the Holm-Šídák post-test. The entire membranes for pro-caspase-1, caspase-1, pro-IL-1β, and IL-1β western blots are shown in Figure S9 and S10.

The incubation of CC endothelial cells with NS13001, a KCa2.2/2.3 opener, or MCC950 (NLRP3 inhibitor) significantly reversed the ET-1-induced reduction of the background membrane conductance (Figure 3c) and reduced the NLRP3 activity (Figure 3d & 3e) and increased K_Ca_2.3 (Figure 3f). Additionally, the incubation with apamin, a K_Ca_2.2/2.3 blocker, did not alter blockage to the total membrane conductance (Figure 3c) and the NLRP3 activity (Figure 3d & 3e) or K_Ca_2.3 expression (Figure 3f) promoted by ET-1 in endothelial cells from murine CC.

Intracavernosal administration of NS13001 rescued ET-1’s reduction in the ICP/MAP responses to submaximal stimuli at 45, 55, and 65 min. Additionally, apamin did not affect ET-1’s impairment of ICP/MAP after its administration (Figure S2b). ET-1 did not change the ICP/MAP responses evoked by maximal stimuli (Figure S2a). Additionally, NS13001 rescued ET-1’s reduction in the ICP/MAP responses to maximal stimuli (Figure S2c).

### Effect of K_Ca_2.2/2.3 opener on ET1-induced erectile dysfunction and NLRP3 activity in DOCA/salt model of hypertension

A selective K_Ca_2.2/2.3 opener NS13001 restored erectile function (Figure 4c) and normalized systolic blood pressure (Figure 4d) in DOCA/salt-treated mice. NS13001 also prevented the increment in the caspase-1 and IL-1β activity in the CC from DOCA/salt mice (Figure 4e and 4f). Additionally, NS13001 prevented the reduction in the K_Ca_2.3 expression (Figure 4g). Moreover, pro-IL-1β expression reduction in the CC from DOCA/salt mice was prevented by NS13001 treatment (Figure S4b). NS13001 impaired electric-field-stimulation-induced relaxation at 2 and 4 Hz (Figure 5c). NS13001 prevented the impaired CC relaxation to ACh and SNP promoted by DOCA/salt (Figure 5d & 5e, Table 1).

**Figure 4.**
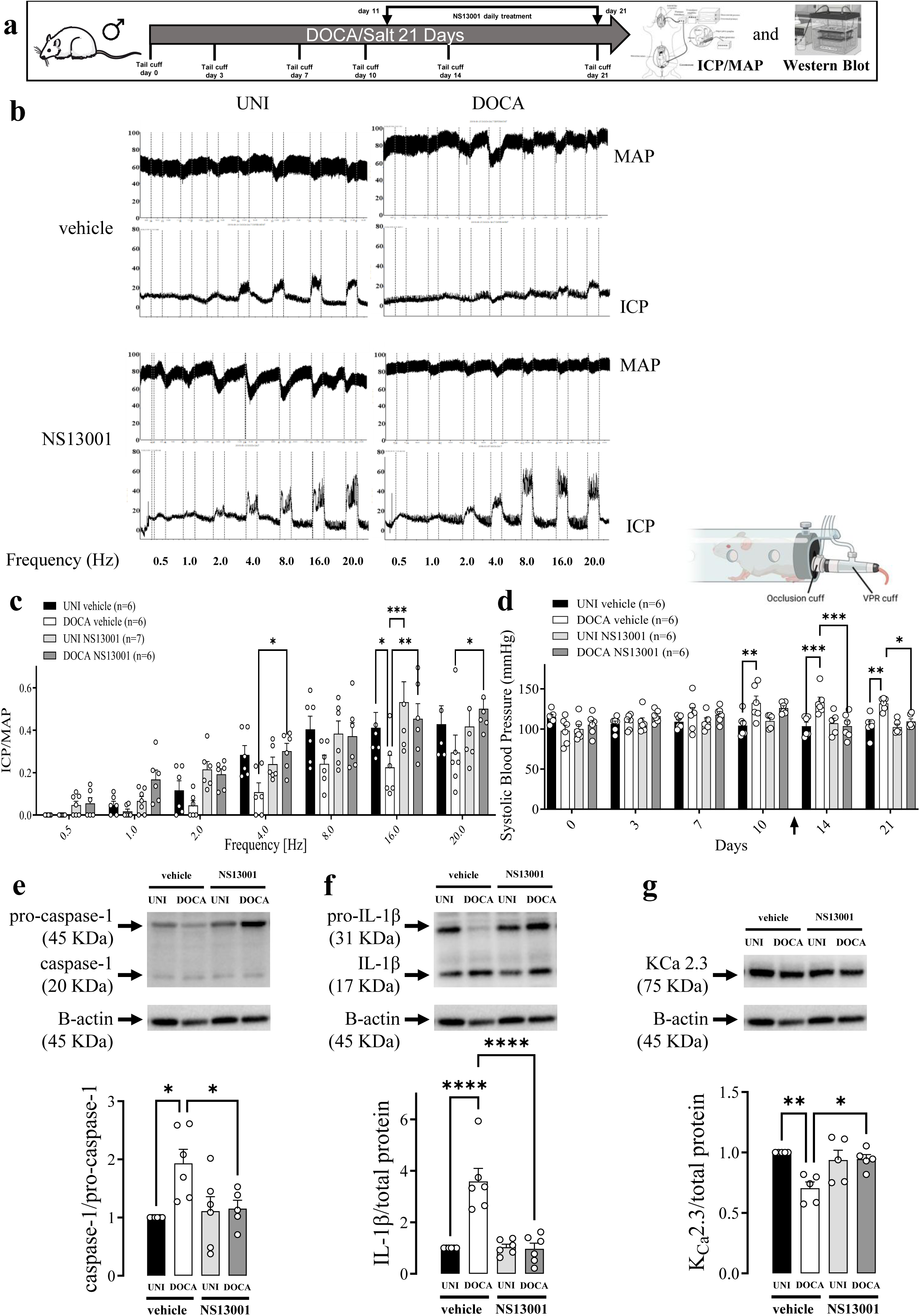
An opener of K_Ca_2 channels, NS13001, restores erectile function and inhibits inflammation in DOCA/salt mice. (**a**) Experimental design. (**b**) Original tracings showing the intracavernosal pressure (ICP, lower traces) and mean arterial pressure (MAP, upper traces) in response to cavernosal nerve stimulation assessed over a range of frequencies (0.5-20 Hz) in unilateral nephrectomized (UNI) and DOCA/salt (DOCA) hypertensive mice treated with vehicle or NS13001. (**c**) Average responses in ICP/MAP and (**d**) systolic blood pressure of UNI and DOCA mice. Analysis of the (**e**) caspase-1, (**f**) IL-1β**, (g)** and K_Ca_2.3 expression in CC strips from DOCA or UNI mice treated with NS13001 or vehicle. Data represent as mean ± SEM values of the groups. * p<0.05. n=5-6. The comparison of each value was performed by two-way ANOVA followed by the Holm-Šídák post-test. ICP=intracavernosal pressure; MAP=mean arterial pressure. The entire membranes for pro-caspase-1, caspase-1, pro-IL-1β, IL-1β, and K_Ca_2.3 western blots are shown in figure S11 and S12.

**Figure 5.**
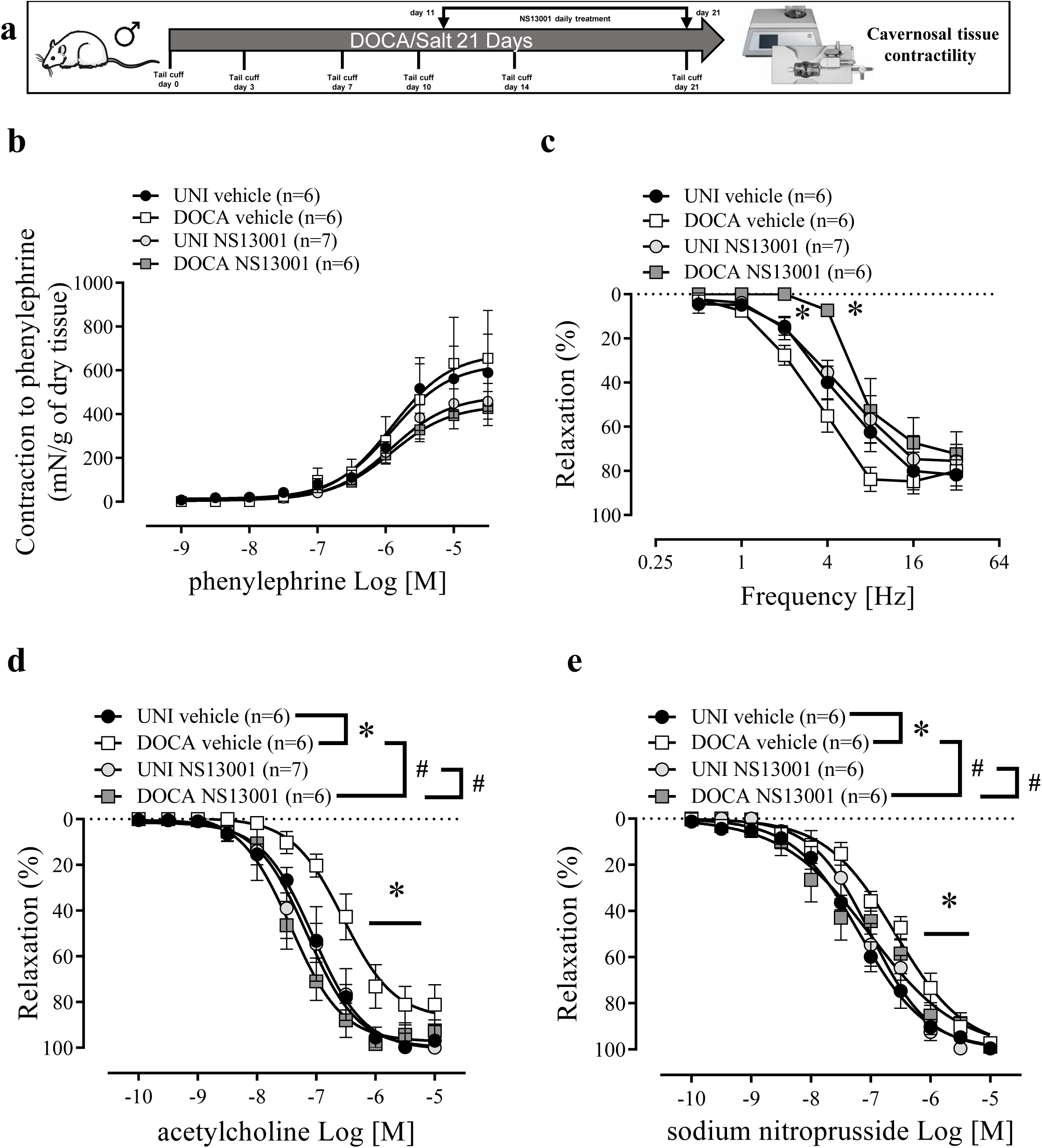
NS13001 restores the impaired endothelium-dependent and -independent relaxations in corpus cavernosum of DOCA/salt mice. (**a**) Experimental design. (**b**) Concentration-response curves for phenylephrine, (**c**) frequency-response curves for electrical-field-stimulation-induced relaxation, and concentration-response curves for (**d**) ACh and (**e**) and SNP in CC strips of DOCA/salt (DOCA) hypertensive and unilaterally nephrectomized (UNI) mice treated with NS13001 or vehicle. The values of the graphics represent as mean ± SEM. * p <0.05. n = 6-8. The comparison of pEC_50_ and Emax (table 1) and each value was performed by two-way ANOVA followed by the Holm-Šídák post-test. Figure 5 (b to e) curves share the same UNI and DOCA vehicles as in Figure 3 (b to e).

### The K_Ca_2.2/2.3 blocker apamin did not affect the ET1-induced erectile dysfunction and NLRP3 activity in DOCA/salt model of hypertension

A selective K_Ca_2.2/2.3 blocker, apamin, impaired erectile function (Figure 6c) and elevated the systolic blood pressure in UNI mice (Figure 6d). Curiously, apamin treatment restored erectile function (Figure 6c) and normalized systolic blood pressure (Figure 6d) in DOCA/salt mice. Apamin also prevented caspase-1 and IL-1β increment in the CC from DOCA/salt mice (Figure 6e and f). Pro-caspase-1 expression was reduced by apamin treatment in the CC of DOCA/salt mice (Figure S4c & S4d). Furthermore, apamin prevented K_Ca_2.3 expression reduction in the CC of DOCA/salt mice (Figure 6g). Apamin impaired electric-field-stimulation-induced relaxation at the frequencies 8, 16 and 20 Hz (Figure 7b, Table 1). Apamin also prevented the impaired relaxation to ACh and SNP promoted by DOCA/salt treatment (Figure 7c & 7d, Table 1).

**Figure 6.**
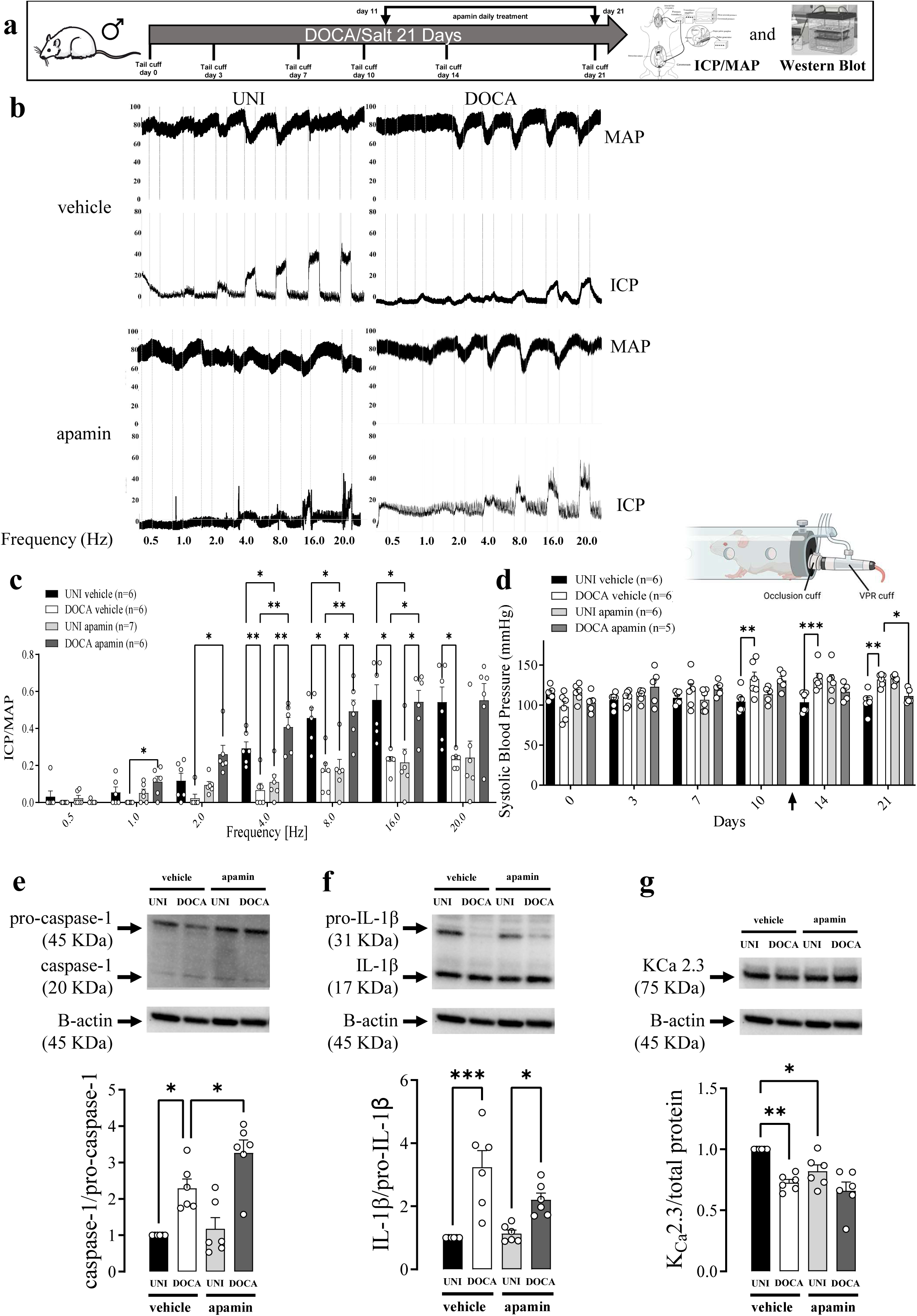
A blocker of K_Ca_2.2/2.3 channels, apamin, inhibits erectile responses in unilaterally nephrectomized control mice and restore responses in DOCA/salt treated mice. (**a**) Experimental design. (**b**) Original tracings showing the intracavernosal pressure (ICP, lower traces) and mean arterial pressure (MAP, upper traces) in response to cavernosal nerve stimulation assessed over a range of frequencies (0.5-20 Hz) in unilateral nephrectomized (UNI) and DOCA/salt (DOCA) hypertensive mice treated with vehicle or apamin. (**c**) Average responses in ICP/MAP and (**d**) systolic blood pressure of UNI and DOCA hypertensive mice. Analysis of the (e) caspase-1, (**f**) IL-1β, and (**g**) K_Ca_2.3 expression in CC strips of DOCA hypertensive or UNI mice treated with apamin or vehicle. Data represent as mean ± SEM values of the groups. * p<0.05. n=5-6. Comparisons of each value were performed using a two-way ANOVA followed by the Holm-Šídák post hoc test. ICP=intracavernosal pressure; MAP=mean arterial pressure. The entire membranes for pro-caspase-1, caspase-1, pro-IL-1β, IL-1β, and K_Ca_2.3 western blots are shown in figure S13 and 14.

**Figure 7.**
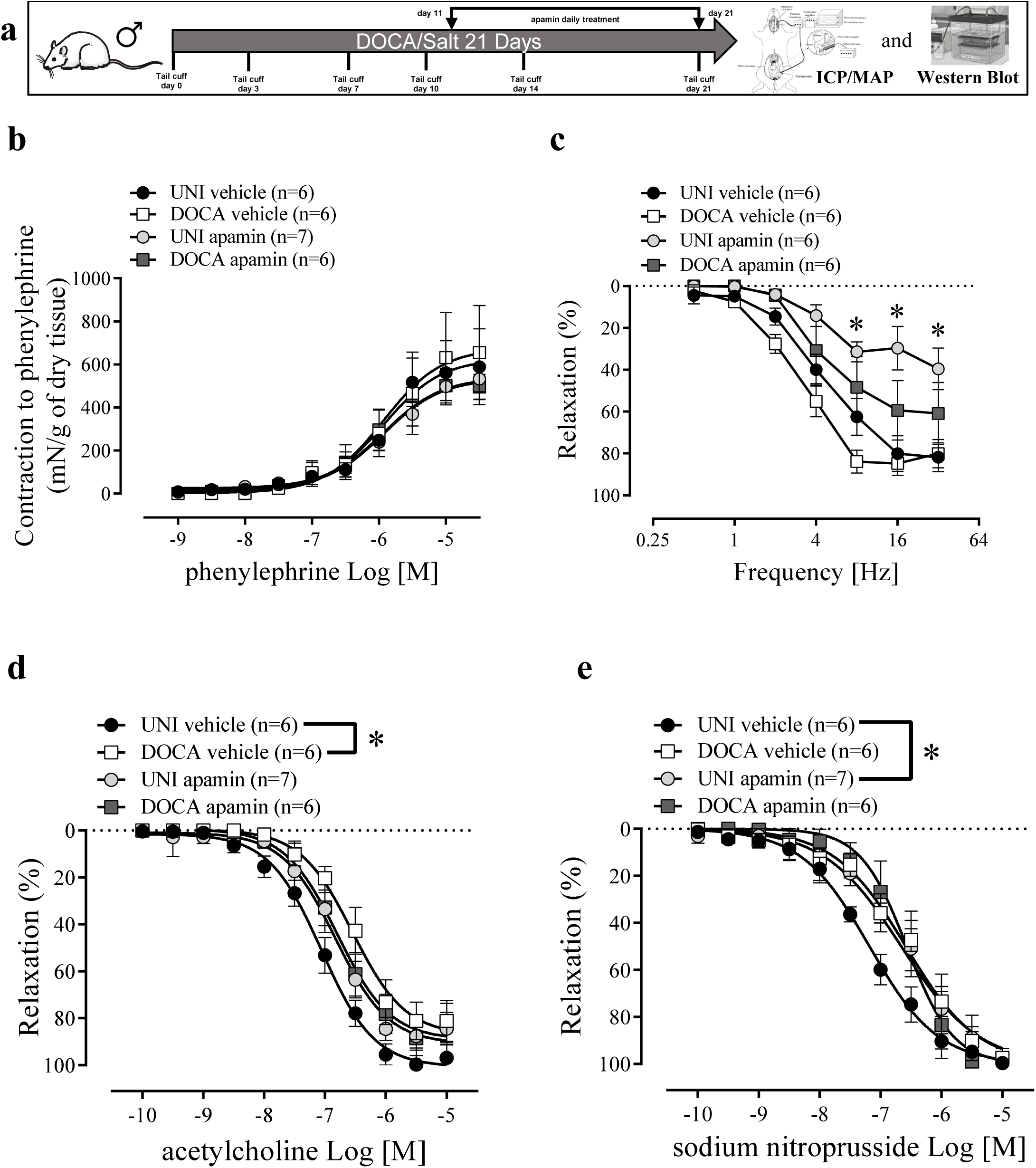
Apamin inhibits relaxation in corpus cavernosum (CC) of unilaterally nephrectomized control mice and has no effect on responses in DOCA/salt mice. (**a**) Experimental design. (**b**) Concentration-response curves for phenylephrine, (**c**) frequency-response curves for electrical-field-stimulation-induced relaxation, and concentration-response curves for (**d**) ACh and (**e**) and SNP in CC strips of DOCA/salt (DOCA) hypertensive and unilaterally nephrectomized (UNI) mice treated with apamin or vehicle. The values of the graphics represent as mean ± SEM. * p <0.05. n = 5-6. The comparison of pEC50 and Emax (table 1) and each value was performed by two-way ANOVA followed by the Holm-Šídák post-test. Figure 8 (a to d) curves share the same UNI and DOCA vehicles as Figure 2 (a to d). Only UNI apamin and DOCA apamin curves were statistically compared.

## 4. Discussion

In this study, we found KCa 2.3 channels play a pivotal role in counteracting ET-1-induced NLRP3 activation and subsequent impairment in vascular and erectile function in DOCA/salt hypertensive mice. This study shows that NS13001, a K_Ca_2.3/K_Ca_2.2 channel opener, has therapeutic potential by inhibiting the ET-1 and NLRP3 inflammatory pathways and restoring erectile function. It has been shown that NS13001 has a higher affinity for K_Ca_2.3, and there is no strong evidence for K_Ca_2.2 modulation on endothelial and erectile dysfunction[2,25]. The increase in K_Ca_2.3 expression in hypertensive mice highlights its relevance in hypertension. Furthermore, as the ET-1 receptor antagonist, bosentan, normalized blood pressure and erectile function in DOCA/salt mice, it underscores the central role of ET-1 in these dysfunctions, and we suggest that it is modulated through the potassium channels.

In the previous study, we showed that NLRP3 modulates erectile function in mice[4]. In addition, it was shown that ET-1-induced erectile dysfunction and pro-inflammatory effect depends on NLRP3 activation[26]. These findings offer valuable insights into the intricate interplay among ET-1, K_Ca_2.2/2.3 channels, and NLRP3 inflammasome, suggesting potential targets for novel therapeutic interventions for inflammatory-related diseases and the restoration of erectile function.

In accordance with our study, several previous studies reported that DOCA/salt treatment leads to hypertension in rodents [7,8,27]. We found that the DOCA/salt mouse model presented a gradual increase in blood pressure, impairment in erectile function, and reduced relaxation to ACh and SNP, i.e., endothelium-dependent and endothelium-independent relaxations, respectively[1,21]. The detrimental effect of DOCA/salt hypertension in rodents on erectile dysfunction was reported previously [21,28]. However, the mechanism behind it was not elucidated. We found in this study that it is associated with ET-1 signaling, leading to changes in the expression of K_Ca_2.3 and the NLRP3 inflammasome.

ET-1 treatment leads to endothelial dysfunction, mainly by reducing the function of various populations of potassium channels, such as inward rectifier (Kir)[28], sensitive ATP (KATP)[29,30], voltage-gated (Kv)[31], and calcium-activated (KCa) [15] potassium channels. Moreover, it was already shown that ET-1 exposure releases pro-inflammatory cytokines, like TNF-α [29] and e IL-1β, in monocytes [30] and smooth muscle cells [31]. Accordingly, in this study, treatment with bosentan, an ET-1 type A (ET_A_) and B (ET_B_) receptor antagonist[32], or MCC950, an NLRP3 inhibitor, restored the K_Ca_2.3 abundance and reduced NLRP3 activity in DOCA/salt mice, suggesting that ET-1 activity in the CC of DOCA/salt mice is mediated via NLRP3 activation[4,26]. Furthermore, treatment with bosentan restored blood pressure, erectile function, and ACh- and SNP-induced relaxation, suggesting that ET-1 signaling contributes to these abnormalities in the DOCA/salt model of hypertension. This is in line with previous reports that administration of ET-1 antagonists reduces the elevation of blood pressure, reverses cardiac hypertrophy and arterial remodeling [16], left ventricular fibrosis [17,18], inflammation, and renal hypertrophy [18].

Accordingly, ET-1 reduces the total membrane conductance in endothelial cells, and our interventions with the K_Ca_2.2/2.3 activator and inhibitor, or NLRP3 inhibitor suggest that this effect is primarily because of the impairment to K_Ca_2.2/2.3 function and NLRP3 inhibition. Notably, the K_Ca_2.3 is expressed explicitly in endothelial but not smooth muscle cells of corpus cavernosum[2]. The K_Ca_2.2 is also expressed in mouse corpus cavernosum, but the role and location of the expression is less known[33] . It has previously been shown that ET-1 leads to endothelial dysfunction in the pulmonary artery, but mainly, this effect was mediated via reduction of the intermediate-conductance calcium-activated potassium channels, i.e., K_Ca_3.1[15]. ET-1 was also shown to inhibit the K_Ca_1.1 in cultured smooth muscle cells from porcine coronary artery [34]. Therefore, this is the first time the inhibitory effect of ET-1 on K_Ca_2.3/2.2 activity has been reported in endothelial cells.

Administration of the K_Ca_2.2/2.3 modulator, NS13001, or MCC950, restored the K_Ca_2.2/2.3 current in endothelial cells incubated with ET-1. Interestingly, it also reduced the NLRP3 activity and restored both endothelium-dependent and endothelium-independent relaxations impaired by exposure to ET-1. K_Ca_2.3 channels were suggested to be involved in endothelium-dependent vasodilatation in human and rat intracavernous penile arteries [12,13]. K_Ca_2.3 activation regulates l-arginine uptake through the y+ transporter system and increased NO release[5], which will counteract the effect of ET-1.

The treatment with NS13001 also reduced caspase-1 and pro-IL-β activity and restored K_Ca_2.3 abundance in DOCA/salt mice CC. It can also be mediated via an anti-inflammatory effect of the K_Ca_2.2/2.3 channels, mainly, because it is known to inhibit the release of proinflammatory cytokines such as IL-6, TNF-α, and the migration of immune system cells[35]. Interestingly, the treatment with apamin reduced K_Ca_2.3 expression in the CC of unilaterally nephrectomized mice with no effect in DOCA/salt hypertensive mice, possibly because it is already downregulated. Following our suggestion about the anti-inflammatory effect of the K_Ca_2.3, apamin administration increased caspase-1 and IL-β activity in DOCA/salt mice CC but not in UNI. Here, it is possible to suggest that the reduction in the K_Ca_2.3 expression correlates with the impairment of its function, once the apamin treatment reduces the K_Ca_2.3 expression. Additionally, apamin has a pro-inflammatory action [36]. Curiously, apamin lowered blood pressure and increased erectile function in DOCA/salt mice. Apamin has been previously shown to protect against atherosclerosis by reducing lesion burden, lipid levels, inflammation, and fibrotic markers[37]. Furthermore, modulating apamin-sensitive SK (small-conductance Ca^2+^-activated K^+^) current, which influences action potential duration and arrhythmia in failing ventricles, selectively reduces nNOS after-hyperpolarization, and its effects on SK channels can be modified by nNOS inhibition [38–40]. Indeed, we observed that apamin increased the CC response to non-adrenergic and non-cholinergic stimulation, suggesting that apamin may be reducing the after-hyperpolarization of the cavernosal nerve.

Importantly, the MCC950, a specific NLRP3 inhibitor, restores a mix of K_Ca_2.3 and other ionic currents. In fact, several Ca^2+^-channels [transient related canonic (TRC) or potential (TRP) channels] or H^+^-channels could also lead to K^+^ efflux from the cell, and this mechanism could be responsible for NLRP3 activation [19]. Then, the MCC950 mechanism of NLRP3 inhibition could be related to the modulation of these channels. Additionally, an administration of MCC950 prevented the erectile dysfunction induced by ET-1. Previous data from our lab had already shown that the NLRP3 genetic deletion or pharmacological inhibition prevents the impairment of endothelium-dependent relaxation evoked by ET-1 in mice CC[26]. Indeed, ET-1, via ET receptor A or B increase cytosolic Ca^2+^ concentration in vascular smooth muscle cells [41], liver cells and cardiomyocytes[42,43].

### Limitations

Limitations to this study include the possibility of investigating only males. This study focused on the vascular mechanisms of erectile dysfunction, which opens the possibility to study only males. In addition, the ICP/MAP ratio was used to measure the erectile function under isoflurane anesthesia. Isoflurane is known to depress the cardiovascular system not only by direct action but also by suppression of the sympathetic nervous system **[44,45]**. This depression in sympathetic activity is closely linked to the increased blood pressure in DOCA/salt mice. In the present study, we cannot exclude off-target effects of the small molecules used to activate K_Ca_2.2/2.3 channels and of the endothelin receptor antagonist, bosentan. In previous studies, we examined the effect of global NLRP3 knockdown in mice, as well as the up- and downregulation of K_Ca_2.3 channels, on erectile function [2,4]. Other genetic approaches allowing tissue-specific knockout and/or genetic manipulation by use of Cas/CRISPR may provide additional information on the effect on the NLRP3 inflammasome by modulation of K_Ca_2.3 channels [46,47].

## 5. Conclusions

In conclusion, our study shows the role of K_Ca_ 2.3 channels in ET-1-induced NLRP3 activation and subsequent impairment in vascular and erectile function in DOCA/salt hypertensive mice. The novel findings highlight the therapeutic potential of the K_Ca_ 2.2/2.3 channel modulator, NS13001, in inhibiting the ET-1 and NLRP3 inflammatory pathways, ultimately leading to the restoration of erectile function. The observed compensatory increase in the K_Ca_2.3 expression in hypertensive mice underscores its significance in hypertension. Furthermore, the efficacy of the ET-1 receptor antagonist, bosentan, in attenuating these effects emphasizes the central role of ET-1 in inducing vascular and erectile dysfunction through the modulation of potassium channels. Our results provide valuable insights into the complex interplay between ET-1, K_Ca_ 2.2/2.3 channels, and NLRP3 inflammatory-related diseases, paving the way for potential therapeutic interventions targeting these pathways to ameliorate inflammatory-related diseases and restore erectile function.

## Abbreviations

The following abbreviations are used in this manuscript:

Ca^2+^: Calcium
K^+^: Potassium
K_Ca_ 2.3: Ca2+-activated K+ channels type 2.3
ET-1: endothelin-1
ICP: intracavernous pressure
CC: corpus cavernosum
IL: interleukin
SK_Ca_: Small conductance Ca^2+^-activated K^+^ channels
IK_Ca_: Intermediate conductance Ca^2+^-activated K^+^ channels
BK_Ca_: Large conductance Ca^2+^-activated K^+^ channels
ED: Erectile dysfunction
NO: Nitric oxide
eNOS: Endothelial NO synthase
TNF-α: Tumor necrosis factor-alpha
K_ir_: Inward-rectifying K^+^ channels
K_ATP_: ATP-sensitive K^+^ channels
K_v_: Voltage-gated K^+^ channels
DOCA: Deoxycorticosterone acetate
UNI: Unilaterally nephrectomized
SBP: Systolic blood pressure
ICAM2: Intracellular adhesion molecule 2
MAP: mean arterial pressure
ACh: Acetylcholine
SNP: Sodium nitroprusside

## Acknowledgments

We want to express profound gratitude for the support from the funding grants PHAB Level 3, AHA Postdoctoral fellowship (24POST1243617), the Sao Paulo Research Foundation (FAPESP, 2016/11988-5; 2016/07641-0; 2018/05638-7; 2019/19749-8); Novo Nordisk Foundation (NFF130C0007739; NFF60C0023284) and Wilhelm Pedersen and Wife’s memorial grant.

