## Supplementary material for "A K_Ca_ 2.2/2.3 Opener Reverses ET-1-induced NLRP3 Activation in Hypertensive Mice Corpora Cavernosa": Figure S

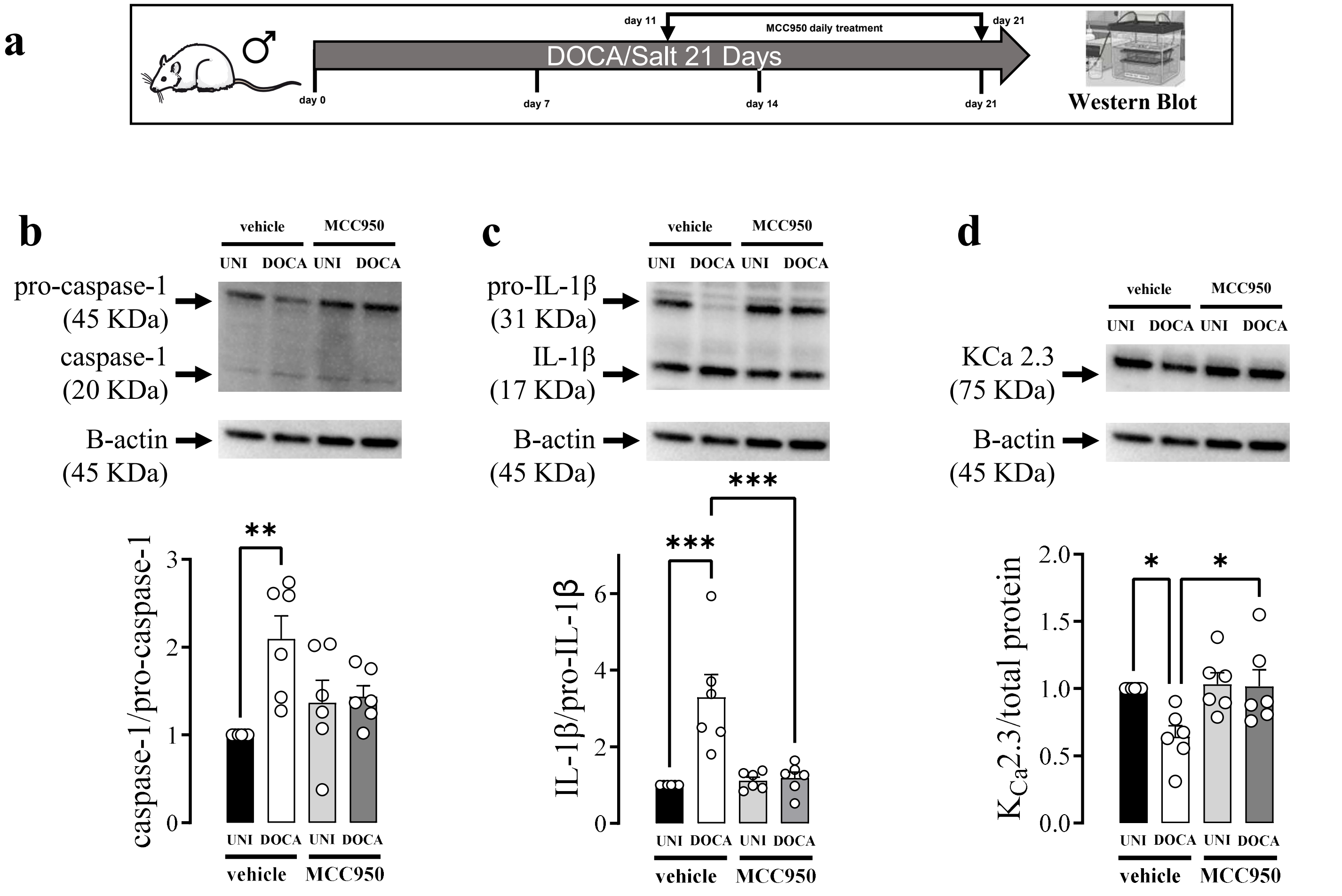


**Figure S1. NLRP3 inhibition prevents inflammation in DOCA/salt mice.** (**a**) Experimental design. Immunoblot analysis of (**b**) caspase-1, (**c**) IL-1β, and **(d)** $K_{Ca}2.3$ expression in CC strips of UNI and DOCA hypertensive mice treated with MCC950 or vehicle. The arrow indicates when the treatment started (c). Data represent the mean ± SEM values of the groups. * $p<0.05$. n=6. The comparison of each value was performed by two-way ANOVA followed by the Holm-Šídák post-test. The entire membranes for pro-caspase-1, caspase-1, pro-IL-1β, IL-1β, and $K_{Ca}2.3$ are shown in figure S7&8.

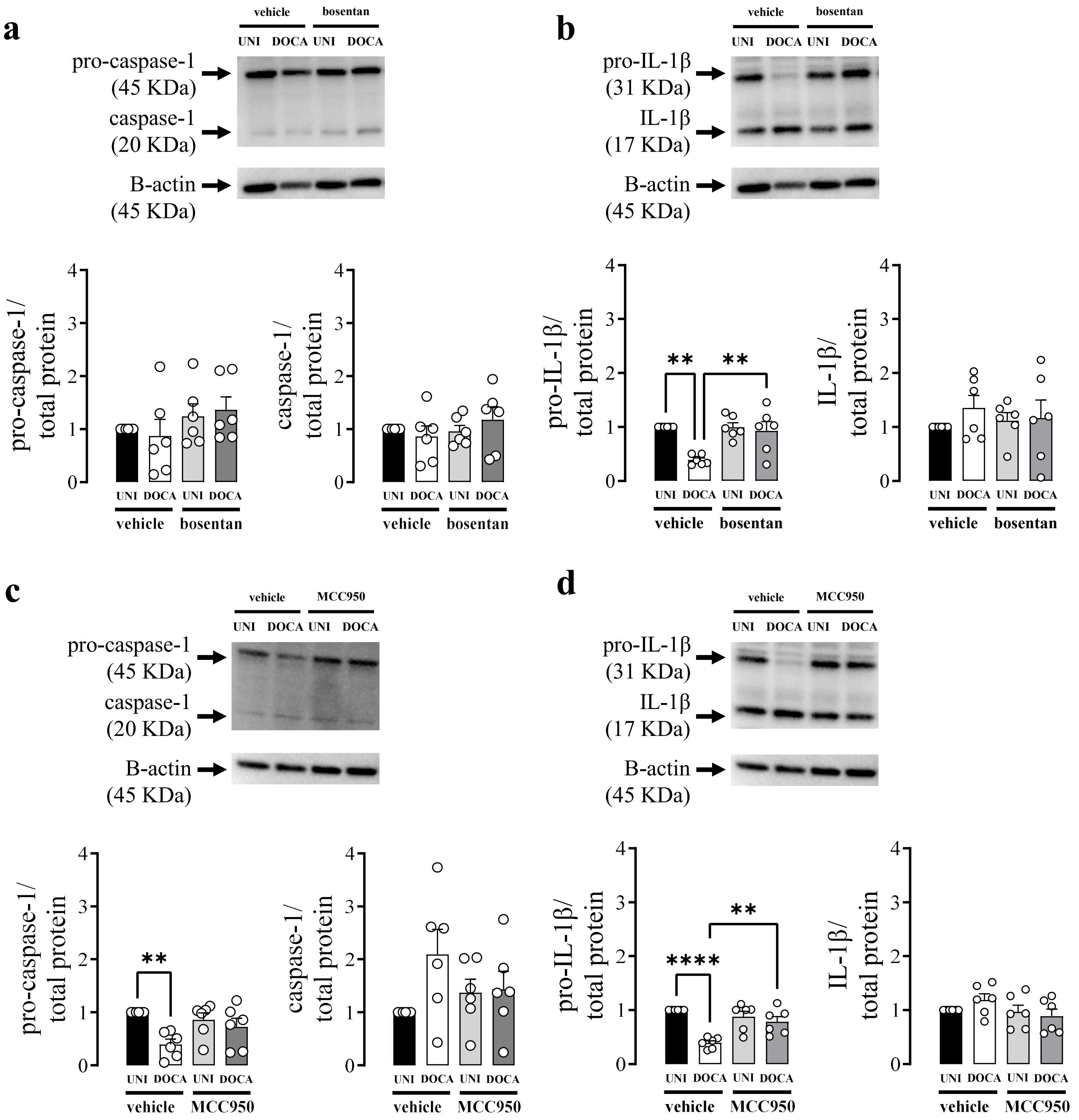


**Figure S2**. Analysis of the (**a** and **c)** pro-caspase-1and caspase-1, (**b** and **d)** pro-IL-1β and IL-1β expression in CC strips of DOCA/salt hypertensive (DOCA) or unilaterally nephrectomized (UNI) rats treated with vehicle, (**a** and **b**) bosentan or (**c** and **d**) MCC950. Data represent the mean ± SEM values of the groups. * p<0.05. n=6. Each value was compared by two-way ANOVA followed by The Holm-Šídák post-test. ICP=intracavernosal pressure; MAP=mean arterial pressure.

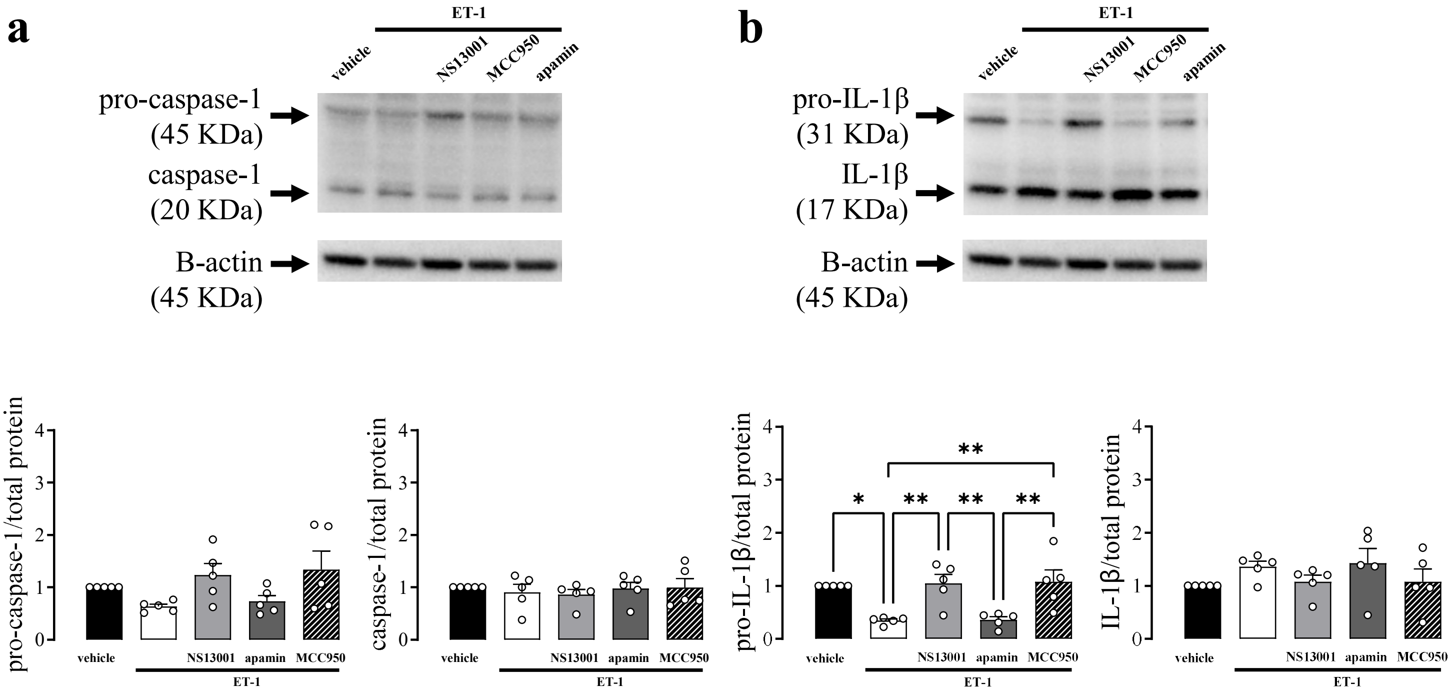


**Figure S3**. Analysis of the (**a**) pro-caspase-1and caspase-1, (**b**) pro-IL-1β and IL-1β expression in CC strips of healthy mice treated with vehicle, or ET-1 in the presence or absence of NS13001, apamin, and MCC950. Data represent the mean ± SEM values of the groups. * $p<0.05$. n=6. Each value was compared by two-way ANOVA followed by The Holm-Šídák post-test. ICP=intracavernosal pressure; MAP=mean arterial pressure.

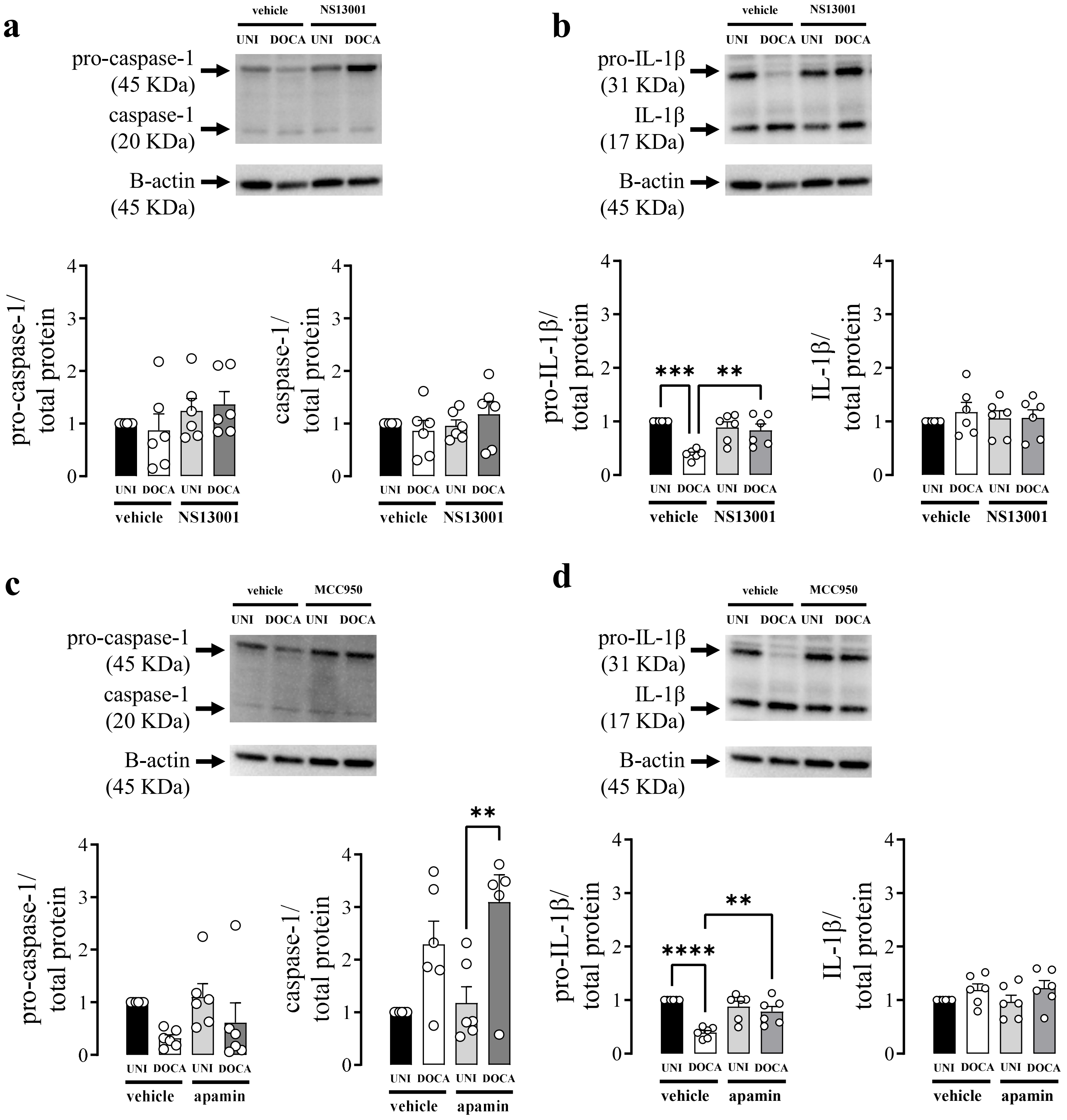


**Figure S4**. Analysis of the (**a** and **c)** pro-caspase-1and caspase-1, (**b** and **d)** pro-IL-1β and IL-1β expression in CC strips of DOCA/salt hypertensive (DOCA) or unilaterally nephrectomized (UNI) mice treated with vehicle, (**a** and **b**) NS13001 or (**c** and **d**) apamin. Data represent the mean ± SEM values of the groups. * p<0.05. n=6. Each value was compared by two-way ANOVA followed by The Holm-Šídák post-test. ICP=intracavernosal pressure; MAP=mean arterial pressure.

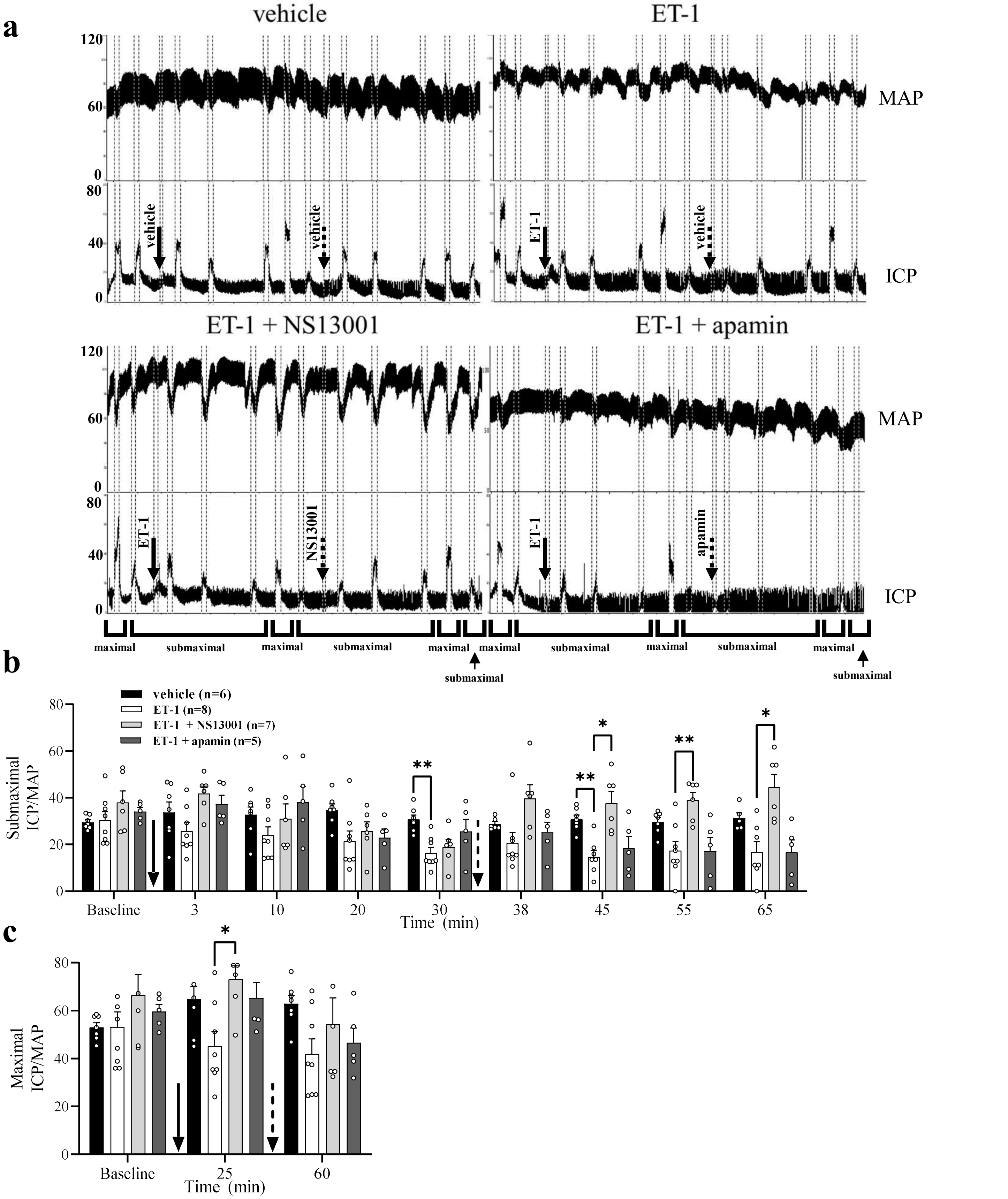


**Figure S5. Effect of $K_{Ca}2.3$ opener or inhibitor on ICP/MAP ratio**. (**a**) Original tracings showing the intracavernosal pressure (ICP, lower traces) and mean arterial pressure (MAP, upper traces) in response to cavernosal nerve stimulation assessed. The graphics depict the ICP/MAP ratio in response to cavernosal nerve stimulation: (**b**) submaximal (2, 4 or 8 Hz) at 3, 10, 20 and 30 min and (**c**) maximal (16 Hz) at 25 min after the injection of the vehicle followed by a new vehicle administration. Submaximal stimulation at 35, 38, 45, 55, and 65 min and maximal at 60 min, ET-1 followed by vehicle, NS13001 or apamin followed by ET-1 or vehicle. Data represent mean ± SEM of ICP/MAP values. Representative tracings showing changes in intracavernosal pressure and blood pressure in response to electrical stimulation of the cavernosal nerve. *p < 0.05. *n* = 5-9. Each frequency value for the ICP/MAP was performed by repeated-measures two-way ANOVA followed by Holm- Šídák post-test. ICP = intracavernosal pressure; MAP = mean arterial pressure. The continuous arrow represents the first administration at 0 min and the dashed arrow represents the second administration at 35 min

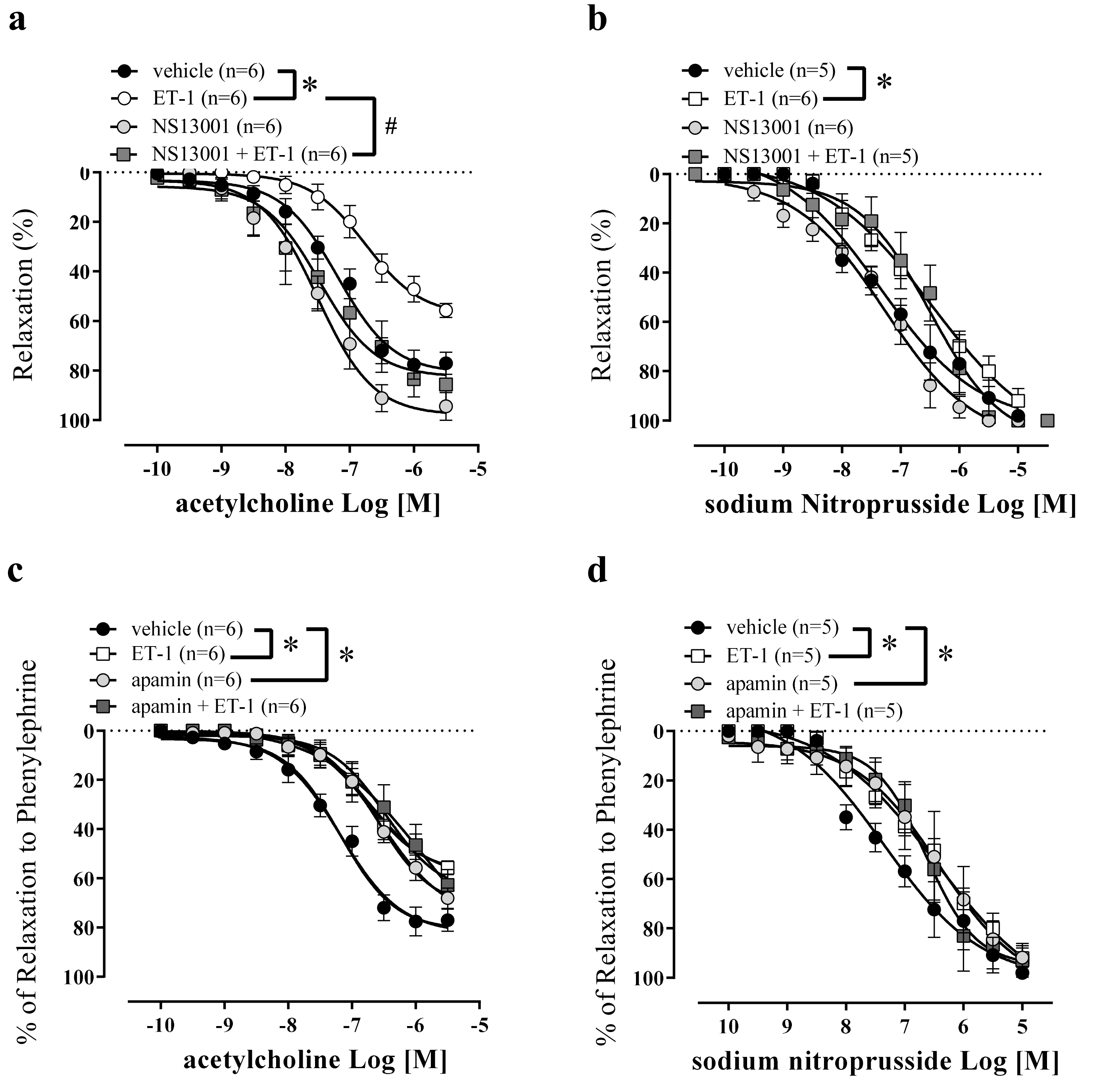


**Figure S6.** Concentration-response curves to (**a** and **c**)acetylcholine, and sodium (**b** and **d**) nitroprusside in CC strips incubated with (**a** and **b**) NS13001 or (**c** and **d**) apamin of healthy mice. * p <0.05. n = 5-6. The comparison of pEC50 and Emax (table 1) and each value was performed by two-way ANOVA followed by Holm-Šídák post-test.

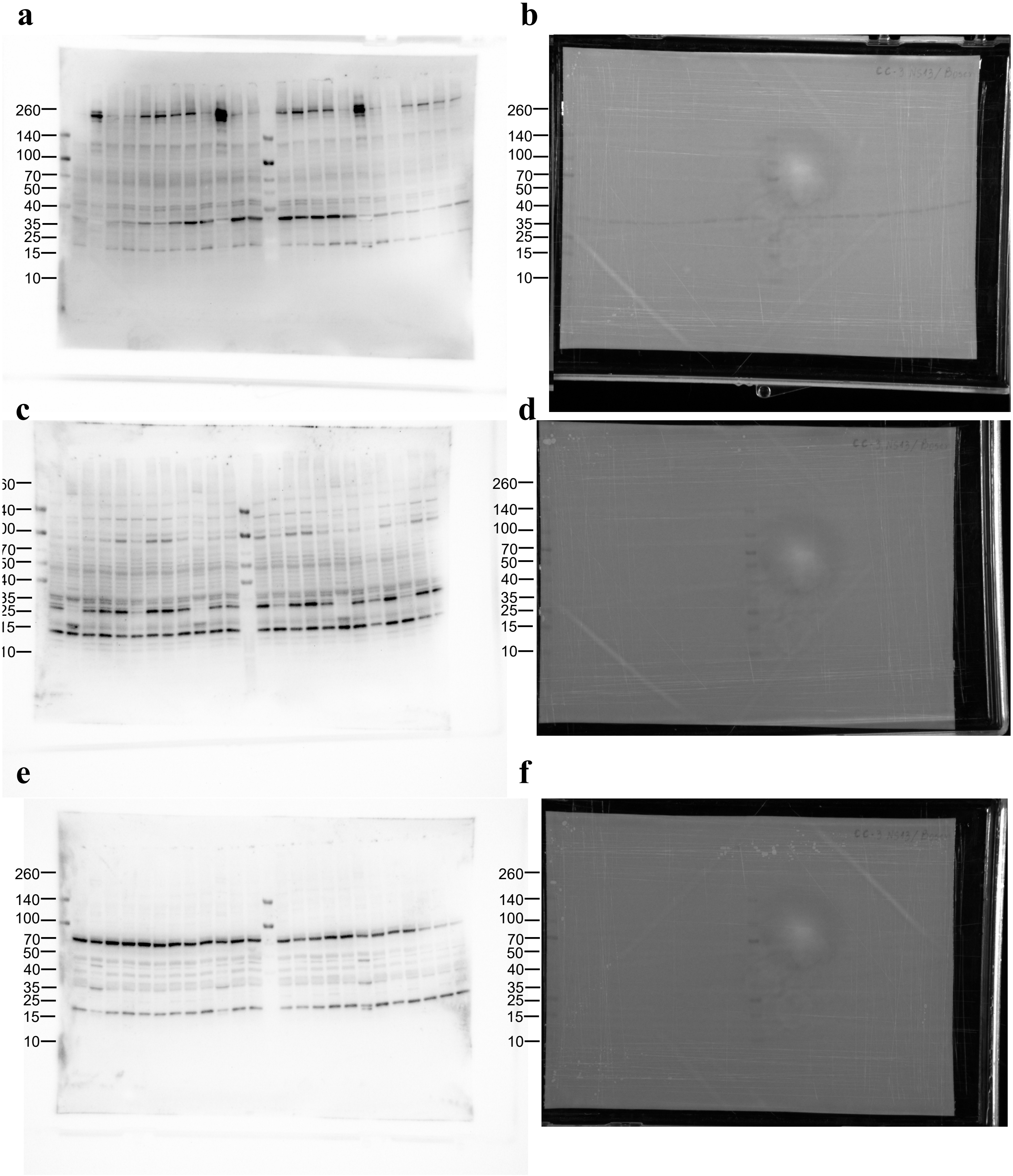


lane 1 – CC UNI vehicle
lane 2 – CC DOCA vehicle
lane 3 – CC UNI bosentan
lane 4 – CC DOCA bosentan
lane 5 – CC UNI vehicle
lane 6 – CC DOCA vehicle
lane 7 – CC UNI bosentan
lane 8 – CC DOCA bosentan

lane 9 – CC UNI vehicle
lane 10 – CC DOCA vehicle
lane 11 – CC UNI bosentan
lane 12 – CC DOCA bosentan
lane 13 – CC UNI vehicle
lane 14 – CC DOCA vehicle
lane 15 – CC UNI NS13001
lane 16 – CC DOCA NS13001

lane 17 – CC UNI vehicle
lane 18 – CC DOCA vehicle
lane 19 – CC UNI NS13001
lane 20 – CC DOCA NS13001
lane 21 – CC UNI vehicle
lane 22 – CC DOCA vehicle
lane 23 – CC UNI NS13001
lane 24 – CC DOCA MCC950

**Figure S7**. The membranes exposed to cheminoluminescence (**a, c, d**) and epiluminescence (**b, d, f**) to develop the immunoblotting for (**a, b**) pro-caspase-1, caspase-1, (**c, d**) pro-IL-1β, IL-1β, (**e, f**) and $K_{Ca}$ 2.3.

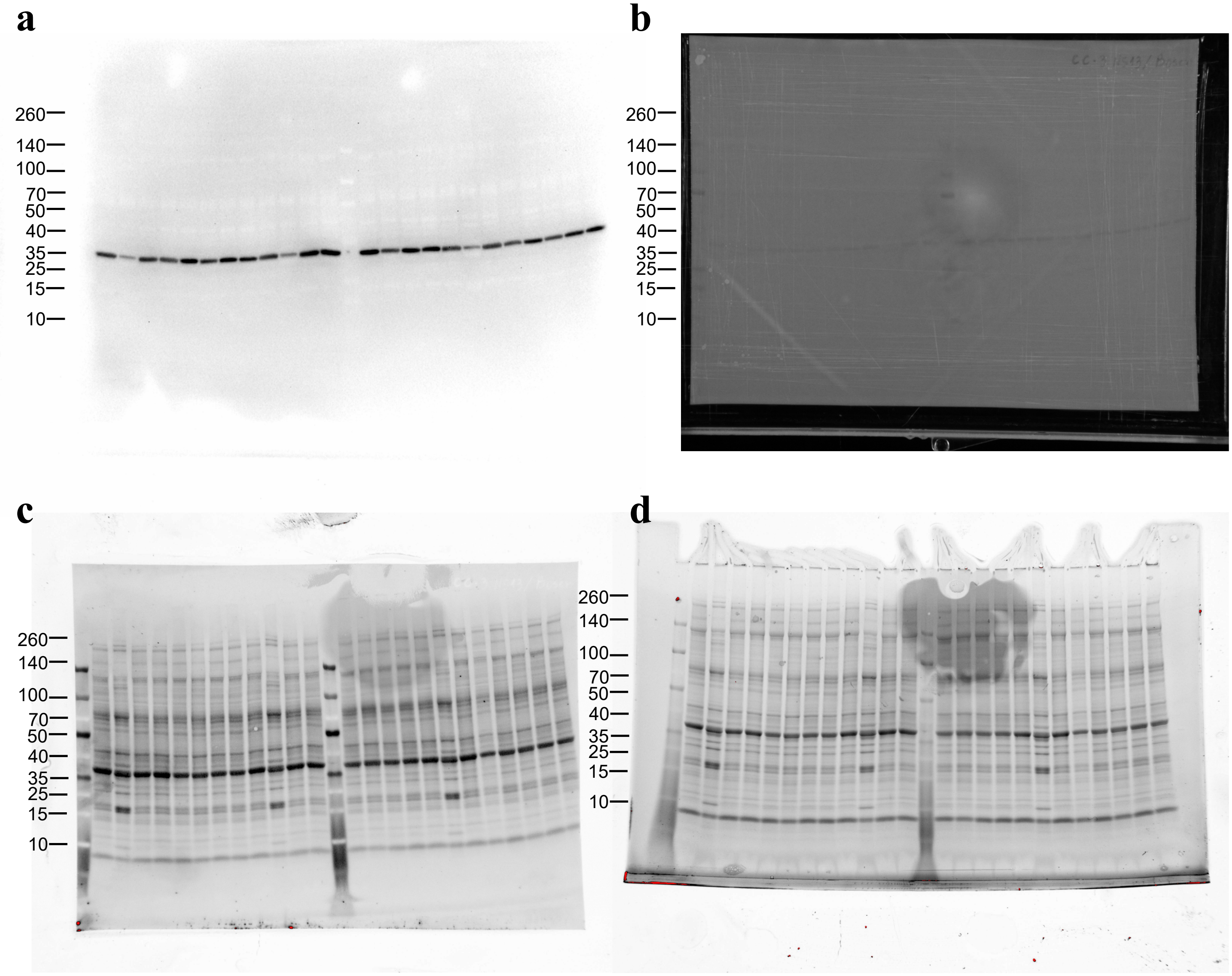


lane 1 – CC UNI vehicle
lane 2 – CC DOCA vehicle
lane 3 – CC UNI bosentan
lane 4 – CC DOCA bosentan
lane 5 – CC UNI vehicle
lane 6 – CC DOCA vehicle
lane 7 – CC UNI bosentan
lane 8 – CC DOCA bosentan

lane 9 – CC UNI vehicle
lane 10 – CC DOCA vehicle
lane 11 – CC UNI bosentan
lane 12 – CC DOCA bosentan
lane 13 – CC UNI vehicle
lane 14 – CC DOCA vehicle
lane 15 – CC UNI NS13001
lane 16 – CC DOCA NS13001

lane 17 – CC UNI vehicle
lane 18 – CC DOCA vehicle
lane 19 – CC UNI NS13001
lane 20 – CC DOCA NS13001
lane 21 – CC UNI vehicle
lane 22 – CC DOCA vehicle
lane 23 – CC UNI NS13001
lane 24 – CC DOCA MCC950

**Figure S8**. The membranes exposed to cheminoluminescence (**a** and **c**) and epiluminescence (**b** and **d**) to develop the immunoblotting for (**a, b**) β-actin. The membranes were exposed to stain-free membrane **(e)** and stain-free gel **(f)** to develop the total protein for membranes.

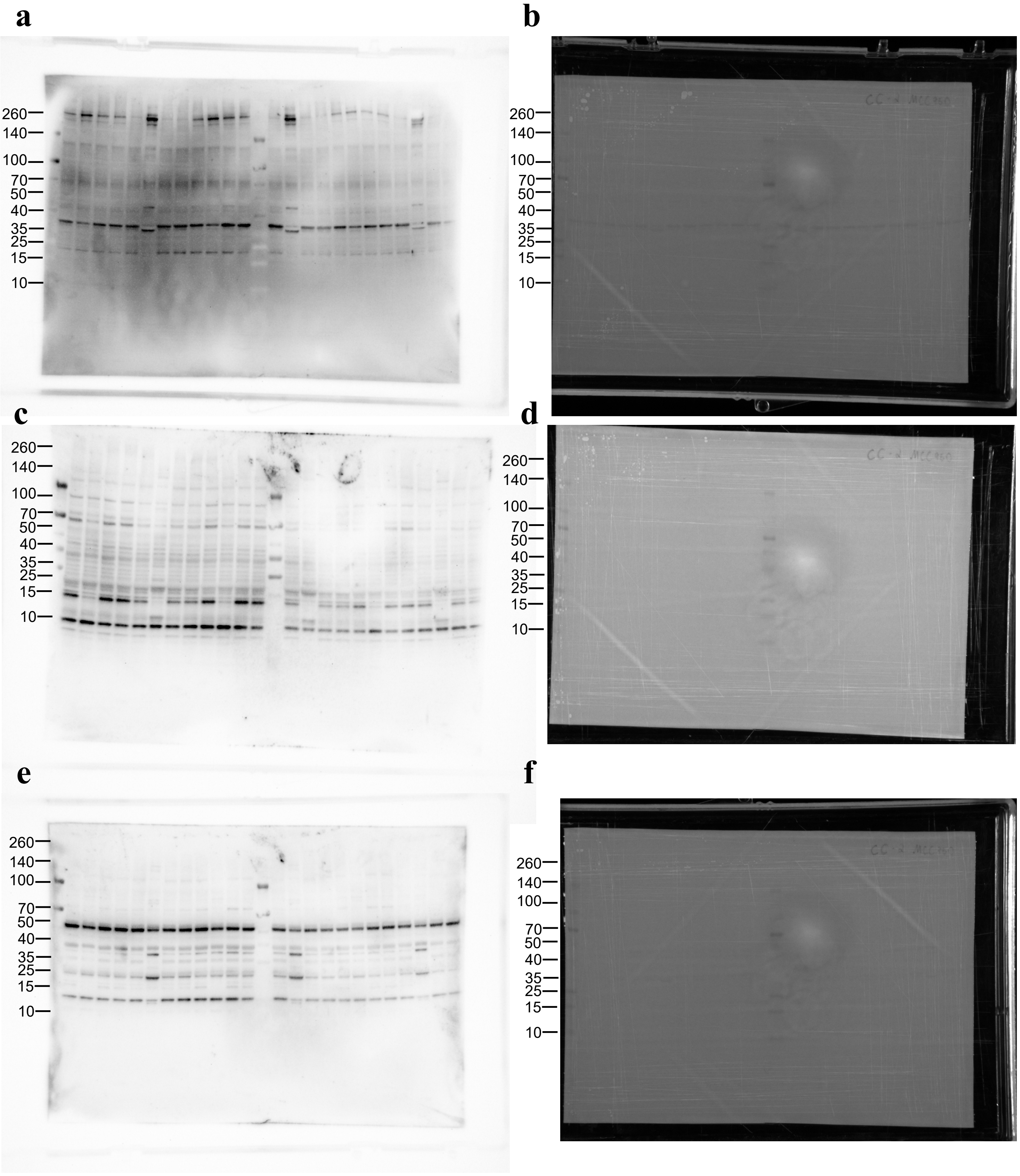


lane 1 – CC UNI vehicle
lane 2 – CC DOCA vehicle
lane 3 – CC UNI MCC950
lane 4 – CC DOCA MCC950
lane 5 – CC UNI vehicle
lane 6 – CC DOCA vehicle
lane 7 – CC UNI MCC950
lane 8 – CC DOCA MCC950

lane 9 – CC UNI vehicle
lane 10 – CC DOCA vehicle
lane 11 – CC UNI MCC950
lane 12 – CC DOCA MCC950
lane 13 – CC UNI vehicle
lane 14 – CC DOCA vehicle
lane 15 – CC UNI MCC950
lane 16 – CC DOCA MCC950

lane 17 – CC UNI vehicle
lane 18 – CC DOCA vehicle
lane 19 – CC UNI MCC950
lane 20 – CC DOCA MCC950
lane 21 – CC UNI vehicle
lane 22 – CC DOCA vehicle
lane 23 – CC UNI MCC950
lane 24 – CC DOCA MCC950

**Figure S9**. The membranes exposed to cheminoluminescence (**a, c, d**) and epiluminescence (**b, d, f**) to develop the immunoblotting for (**a, b**) pro-caspase-1, caspase-1, (**c, d**) pro-IL-1β, IL-1β, (**e, f**) and $K_{Ca}$ 2.3.

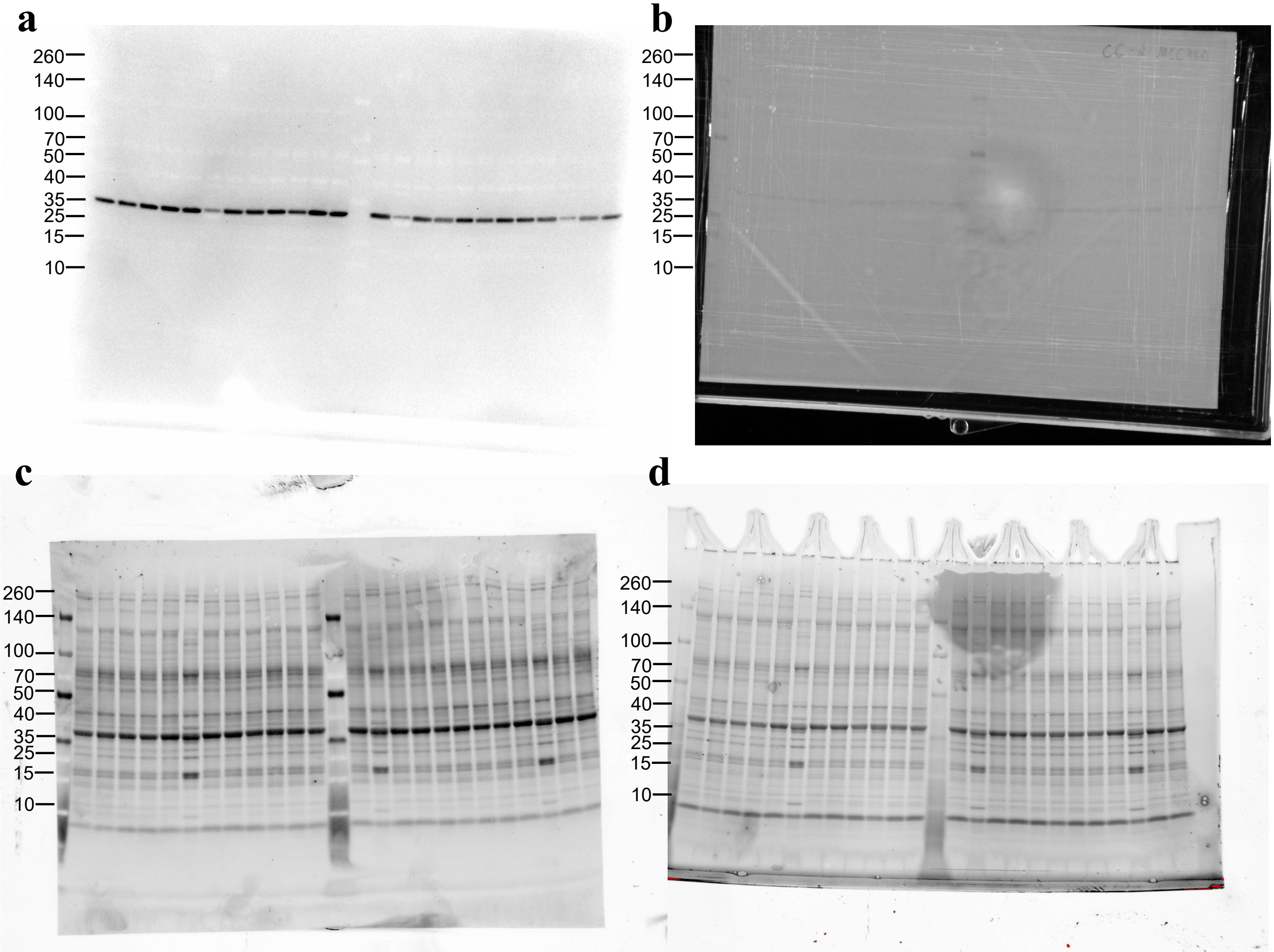


lane 1 – CC UNI vehicle
lane 2 – CC DOCA vehicle
lane 3 – CC UNI MCC950
lane 4 – CC DOCA MCC950
lane 5 – CC UNI vehicle
lane 6 – CC DOCA vehicle
lane 7 – CC UNI MCC950
lane 8 – CC DOCA MCC950

lane 9 – CC UNI vehicle
lane 10 – CC DOCA vehicle
lane 11 – CC UNI MCC950
lane 12 – CC DOCA MCC950
lane 13 – CC UNI vehicle
lane 14 – CC DOCA vehicle
lane 15 – CC UNI MCC950
lane 16 – CC DOCA MCC950

lane 17 – CC UNI vehicle
lane 18 – CC DOCA vehicle
lane 19 – CC UNI MCC950
lane 20 – CC DOCA MCC950
lane 21 – CC UNI vehicle
lane 22 – CC DOCA vehicle
lane 23 – CC UNI MCC950
lane 24 – CC DOCA MCC950

**Figure S10**. The membranes exposed to cheminoluminescence (**a** and **c**) and epiluminescence (**b** and **d**) to develop the immunoblotting for (**a, b**) β-actin. The membranes were exposed to stain-free membrane **(e)** and stain-free gel **(f)** to develop the total protein for membranes.

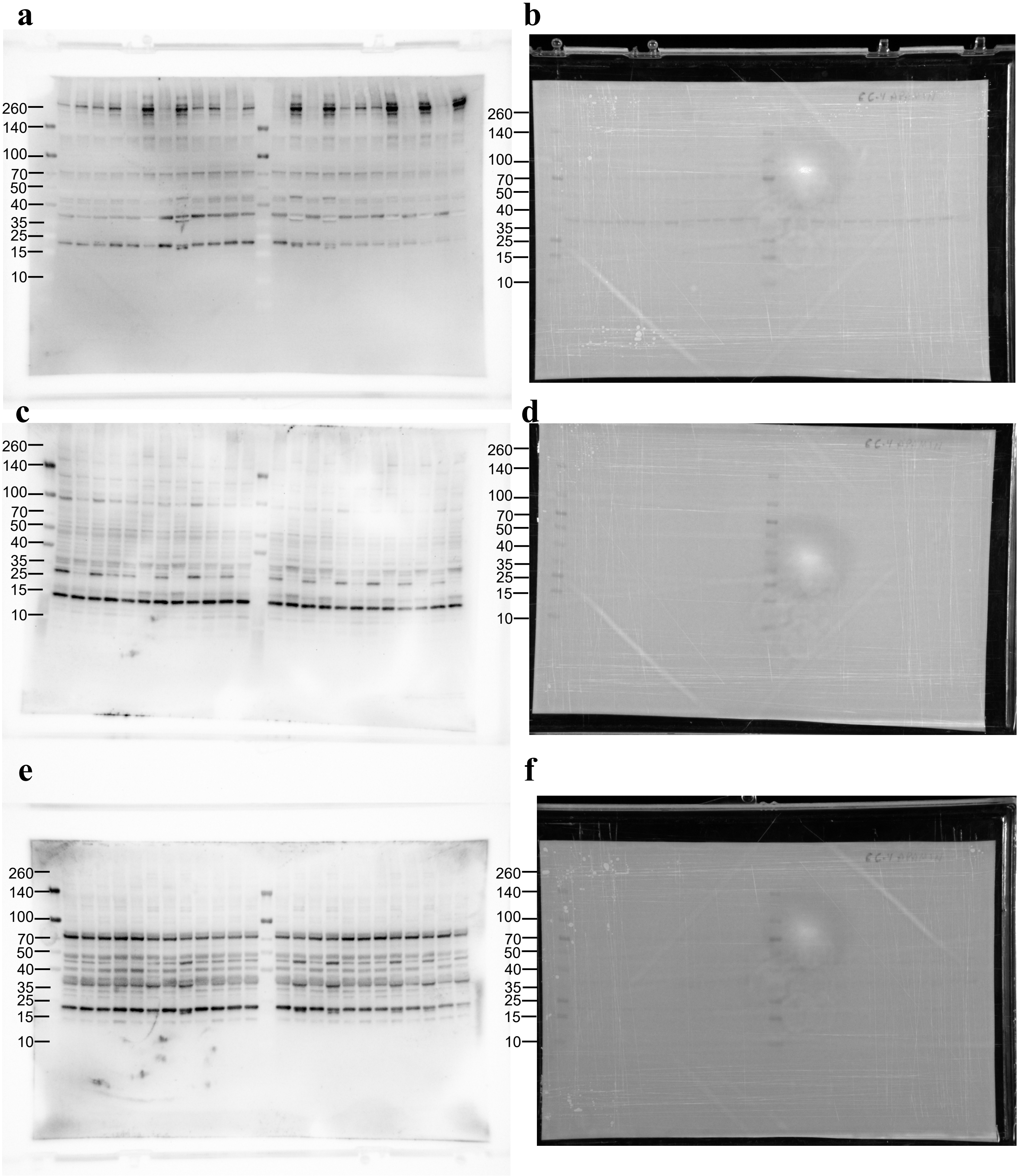


lane 1 – CC UNI vehicle
lane 2 – CC DOCA vehicle
lane 3 – CC UNI apamin
lane 4 – CC DOCA apamin
lane 5 – CC UNI vehicle
lane 6 – CC DOCA vehicle
lane 7 – CC UNI apamin
lane 8 – CC DOCA apamin

lane 9 – CC UNI vehicle
lane 10 – CC DOCA vehicle
lane 11 – CC UNI apamin
lane 12 – CC DOCA apamin
lane 13 – CC UNI vehicle
lane 14 – CC DOCA vehicle
lane 15 – CC UNI apamin
lane 16 – CC DOCA apamin

lane 17 – CC UNI vehicle
lane 18 – CC DOCA vehicle
lane 19 – CC UNI apamin
lane 20 – CC DOCA apamin
lane 21 – CC UNI vehicle
lane 22 – CC DOCA vehicle
lane 23 – CC UNI apamin
lane 24 – CC DOCA apamin

**Figure S11**. The membranes exposed to cheminoluminescence (**a, c, d**) and epiluminescence (**b, d, f**) to develop the immunoblotting for (**a, b**) pro-caspase-1, caspase-1, (**c, d**) pro-IL-1β, IL-1β, (**e, f**) and $K_{Ca}$ 2.3.

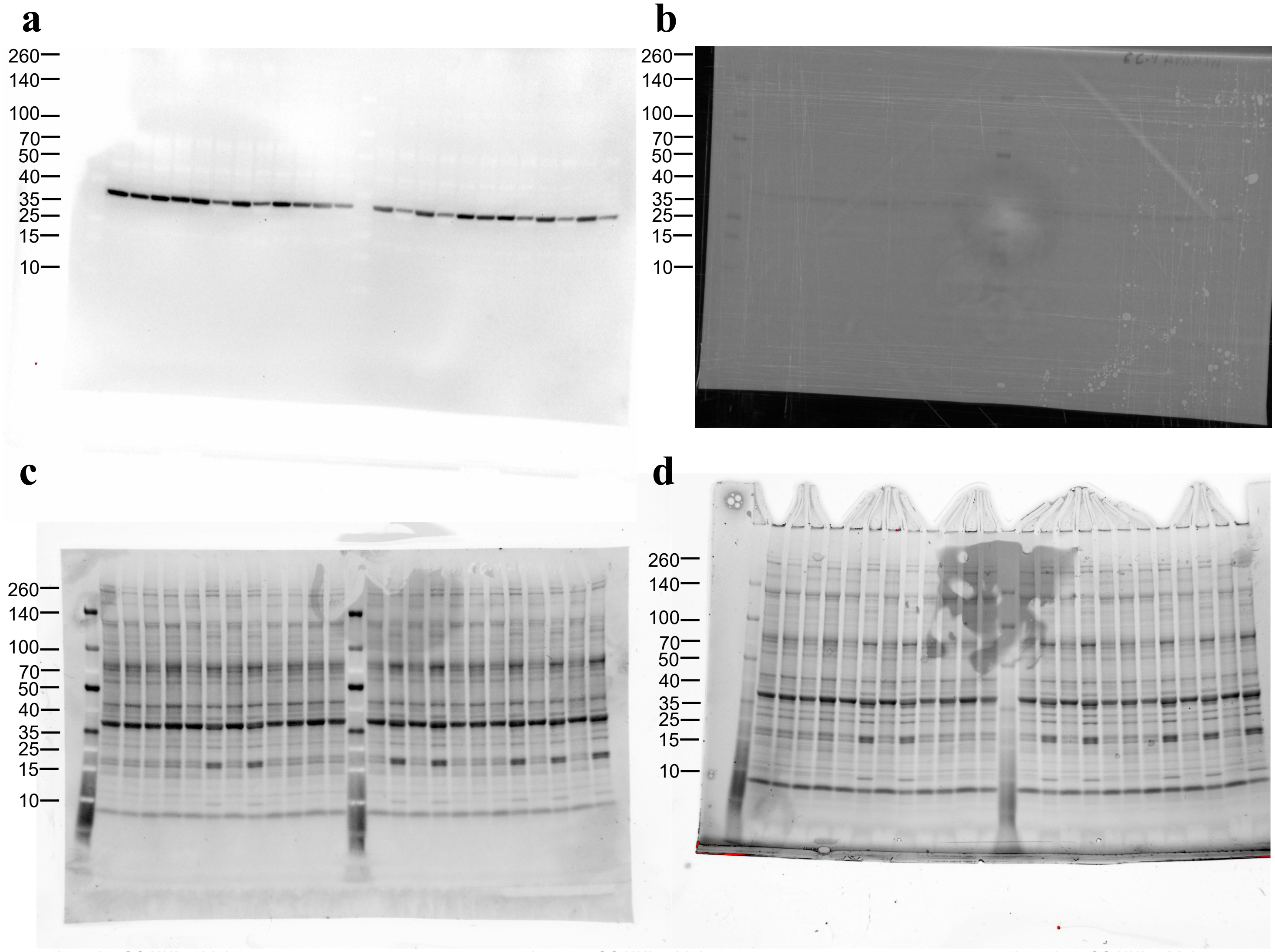


lane 1 – CC UNI vehicle
lane 2 – CC DOCA vehicle
lane 3 – CC UNI apamin
lane 4 – CC DOCA apamin
lane 5 – CC UNI vehicle
lane 6 – CC DOCA vehicle
lane 7 – CC UNI apamin
lane 8 – CC DOCA apamin

lane 9 – CC UNI vehicle
lane 10 – CC DOCA vehicle
lane 11 – CC UNI apamin
lane 12 – CC DOCA apamin
lane 13 – CC UNI vehicle
lane 14 – CC DOCA vehicle
lane 15 – CC UNI apamin
lane 16 – CC DOCA apamin

lane 17 – CC UNI vehicle
lane 18 – CC DOCA vehicle
lane 19 – CC UNI apamin
lane 20 – CC DOCA apamin
lane 21 – CC UNI vehicle
lane 22 – CC DOCA vehicle
lane 23 – CC UNI apamin
lane 24 – CC DOCA apamin

**Figure S12**. The membranes exposed to cheminoluminescence (**a** and **c**) and epiluminescence (**b** and **d**) to develop the immunoblotting for (**a, b**) β-actin. The membranes were exposed to stain-free membrane **(e)** and stain-free gel **(f)** to develop the total protein for membranes.

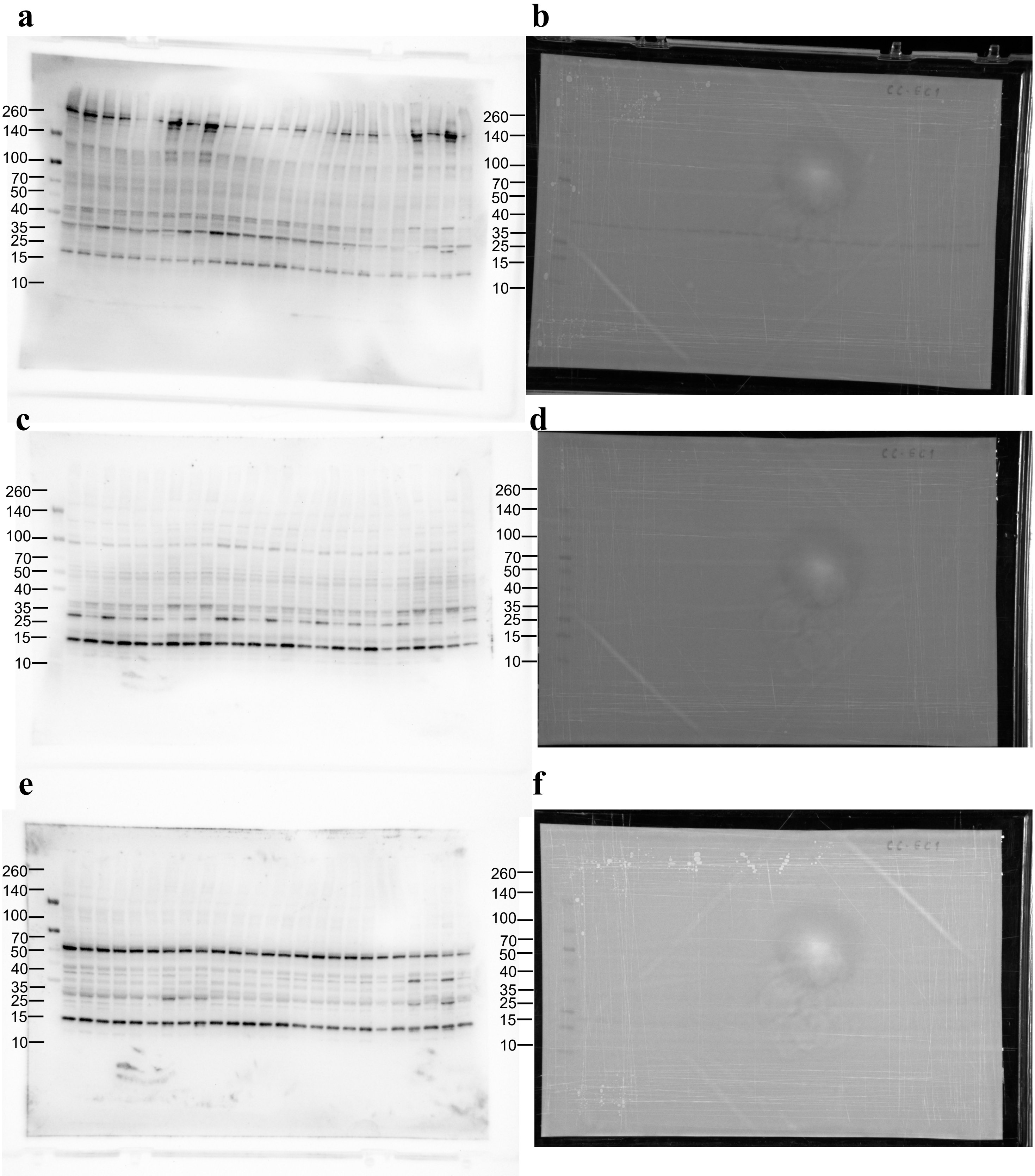


lane 1 – CC vehicle
lane 2 – CC ET-1
lane 3 – CC ET-1 NS13001
lane 4 – CC ET-1 apamin
lane 5 – CC ET-1 MCC950
lane 6 – CC vehicle
lane 7 – CC ET-1
lane 8 – CC ET-1 NS13001
lane 9 – CC ET-1 apamin
lane 10 – CC ET-1 MCC950

lane 11 – CC vehicle
lane 12 – CC ET-1
lane 13 – CC ET-1 NS13001
lane 14 – CC ET-1 apamin
lane 15 – CC ET-1 MCC950
lane 16 – CC vehicle
lane 17 – CC ET-1
lane 18 – CC ET-1 NS13001
lane 19 – CC ET-1 apamin
lane 20 – CC ET-1 MCC950

lane 21 – CC vehicle
lane 22 – CC ET-1
lane 23 – CC ET-1 NS13001
lane 24 – CC ET-1 apamin
lane 25 – CC ET-1 MCC950

**Figure S13**. The membranes exposed to cheminoluminescence (**a, c, d**) and epiluminescence (**b, d, f**) to develop the immunoblotting for (**a, b**) pro-caspase-1, caspase-1, (**c, d**) pro-IL-1β, IL-1β, (**e, f**) and $K_{Ca}$ 2.3.

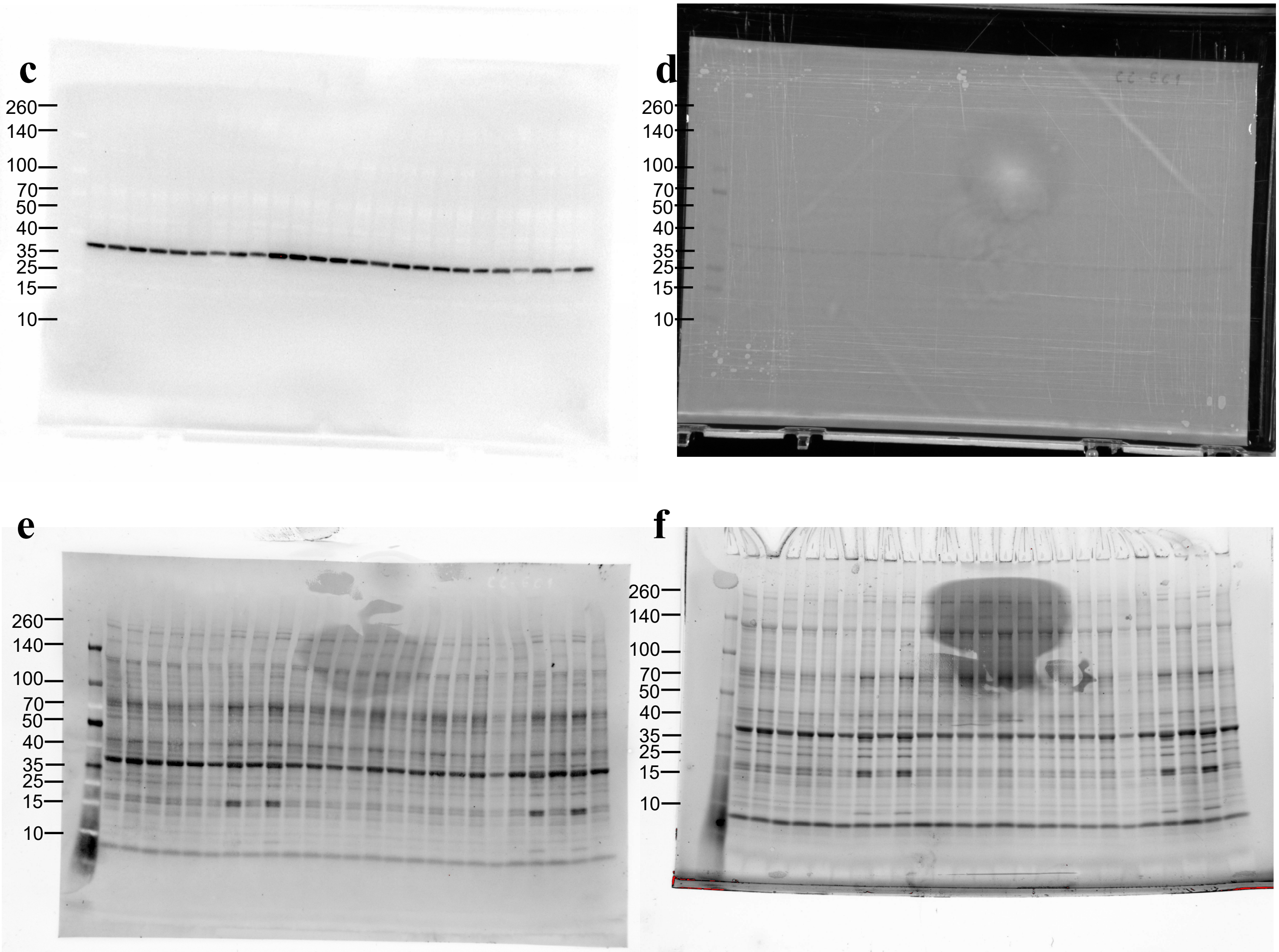


**lane 1 – CC vehicle**
**lane 2 – CC ET-1**
**lane 3 – CC ET-1 NS13001**
**lane 4 – CC ET-1 apamin**
**lane 5 – CC ET-1 MCC950**
**lane 6 – CC vehicle**
**lane 7 – CC ET-1**
**lane 8 – CC ET-1 NS13001**
**lane 9 – CC ET-1 apamin**
**lane 10 – CC ET-1 MCC950**

**lane 11 – CC vehicle**
**lane 12 – CC ET-1**
**lane 13 – CC ET-1 NS13001**
**lane 14 – CC ET-1 apamin**
**lane 15 – CC ET-1 MCC950**
**lane 16 – CC vehicle**
**lane 17 – CC ET-1**
**lane 18 – CC ET-1 NS13001**
**lane 19 – CC ET-1 apamin**
**lane 20 – CC ET-1 MCC950**

**lane 21 – CC vehicle**
**lane 22 – CC ET-1**
**lane 23 – CC ET-1 NS13001**
**lane 24 – CC ET-1 apamin**
**lane 25 – CC ET-1 MCC950**

**Figure S14**. The membranes exposed to cheminoluminescence (**a** and **c**) and epiluminescence (**b** and **d**) to develop the immunoblotting for (**a, b**) β-actin. The membranes were exposed to stain-free membrane **(e)** and stain-free gel **(f)** to develop the total protein for membranes.
